# Electric field mediated fibronectin-hydroxyapatite interaction: A molecular insight

**DOI:** 10.1101/2020.07.22.215327

**Authors:** Subhadip Basu, Biswajit Gorai, Bikramjit Basu, Prabal K. Maiti

**Author notes:** Joint corresponding authors; (B.Basu); (P.K. Maiti).

## Abstract

In experimental research driven biomaterials science, the influence of different material properties (elastic stiffness, surface energy, etc.), and to a relatively lesser extent, the biophysical stimulation (electric/magnetic) on the cell-material interaction has been extensively investigated. Considering the central importance of the protein adsorption on cell-material interaction, the role of physiochemical factors on the protein adsorption is also probed. Despite its significance, the quantitative analysis of many such aspects remains largely unexplored in biomaterials science. In recent studies, the critical role of electric field stimulation towards modulation of cell functionality on implantable biomaterials has been experimentally demonstrated. Given this background, we investigated the influence of external electric field stimulation (upto 1.00 V/nm) on fibronectin (FN) adsorption on hydroxyapatite, HA (100) surface at 300K using all-atom MD simulation method. Fibronectin adsorption was found to be governed by the attractive electrostatic interaction, which changed with the electric field strength. Non-monotonous changes in structural integrity of fibronectin were recorded with the change in field strength and direction. This can be attributed to the spatial rearrangement of local charges and global structural changes of the protein. The dipole moment vectors of fibronectin, water and HA quantitatively exhibited similar pattern of orienting themselves parallel to the field direction, with field strength dependent increase in their magnitudes. No significant change has been recorded for radial distribution function of water surrounding fibronectin. Field dependent variation in the salt bridge nets and number of hydrogen bonds between fibronectin and hydroxyapatite were also examined. One of the important results in the context of the cell-material interaction is that the RGD sequence of FN was exposed to solvent side, when the field was applied along a direction outward perpendicular to HA (001) surface. Summarizing, the present study provides quantitative insights into the influence of electric field stimulation on biomolecular interactions involved in fibronectin adsorption on hydroxyapatite surface.

## 1. Introduction

A number of unmet clinical needs have triggered significant research activities to develop new biomaterials and bioengineering strategies. The quantitative understanding of the nature of interactions between inorganic surfaces and the components of living systems (proteins, cells, bacteria, blood, etc.) is of utmost significance for rational designing of new biomaterials. The very first step of this interaction is the adsorption of proteins on the biomaterial surface. When a biomaterial is implanted in a human subject, the abundance of adsorbed proteins on the material surface initiates a cascade of biophysical events, like platelet activation and foreign body response.^1^ It is the adsorbed proteins, which interact with the cell surface receptors to result in the formation of focal adhesion complexes.^2^ Therefore, the cell-material interaction and cellular functionality changes largely depend on the nature and structural integrity of the adsorbed proteins. Mechanical strength of the natural hard tissue also results from the attachment of collagen and other proteins to its mineral part.^3^ Bacterial attachment on the implant surface is also governed by the adsorbed protein-bacterial protein interaction. Due to such plethora of biophysical implications, the protein-material interaction has acquired the central importance in the field of biomaterials science.

Several physiochemical factors play important roles in protein adsorption phenomena. Protein adsorption kinetics is largely influenced by surface roughness, porosity, particle size, wettability, chemical groups on the substrate, surface defects and vacancies.^4^ The adsorption probability also depends on the size of the protein and its amino acid sequence.^4^ In general, larger protein molecules show a greater potential to be adsorbed on a material substrate. Due to greater diffusion rate, smaller molecules get adsorbed onto the surface first and later are replaced by larger proteins.^5^ Various factors, like pH, and presence of different ions also influence the adsorption process.^4^ Besides these, the protein adsorption phenomena can also be modulated with the application of external stimuli (thermal, electric and magnetic).^6^

A number of computational studies are reported to understand the interaction between HA surface and proteins/peptides/amino acids.^7, 8, 9, 10^ It is now evident that the adsorption of amino acids on the HA surface takes place via formation of Ca-O and H-bonds.^10, 11^ Apart from these, electrostatic interactions among oppositely charged groups (COO^-^ and Ca^2+^, NH_3_^+^ and PO_4_ ^3-^) also regulate the overall adsorption process.^11^ It is quite natural that similar kind of interaction has been observed between whole protein molecules and different HA crystal facets. The adsorption of bone morphogenic protein-2 (BMP-2), for example, has been found to be mediated by the electrostatic interaction between -COO^-^ and Ca^2+^.^12^ Water bridged H-bonds were present between protein and material surface, as well.^12^ The adsorption mechanism of BMP-7, on the other side, was found to be dominated by electrostatic interaction between Ca^2+^ and -COO^-^, together with H-bond formation between NH_3_^+^and PO_4_^3-^.^13^Another important protein is fibronectin (FN), which is an extracellular matrix protein having molecular weight of ∼440 kDa.^14^ Three types of FN repeats (FNI, FNII and FNIII) are found in each monomer of this dimer protein.^14^ Almost 90% of FN sequence is comprised of 12 FNI repeats, 2 FNII repeats and 15-17 FNIII repeats.^14^ The adsorption study of FNIII_10_ module on HA (001) surface established electrostatic interaction force to be the governing factor.^15^ In case of the FNIII_7-10_ module, two stages of adsorption, namely weak adsorption and strong adsorption, were recorded.^16^ While the weak adsorption was driven by VDW interaction energy; coulombic interaction dominated the strong adsorption phase.^16^ The nature of interaction between different proteins like osteopontin (OPN), bone sialoprotein (BSP) and leucine-rich amelogenin protein (LRAP) with the HA surface has also been investigated.^17, 18, 19^

In spite of these existing reports, the effects of external stimuli (electric or magnetic) on protein adsorption on the HA surface remains unexplored. Among various kinds of stimuli, the effect of external electric field on cell functionality modulation has been recently investigated in some of our previous studies.^20, 21, 22^ Although the beneficial effect of electric field stimulation on bone cell growth and bone healing is relatively well-established, little is known about the underlying mechanisms in the context of protein-material interaction.^23^ Against this background, the present study aims to bridge this gap by exploring the effect of electric field stimulation on protein-HA interaction, which, in turn, mediates cell-material interaction. The 10^th^ type III module of fibronectin has been chosen as model protein and the external electric field dependent protein conformational changes have been probed quantitatively. The changes in dipole moment orientation, hydrogen bond formation and interaction energies with the strength and direction of electric field have been assessed. In summary, the present study establishes quantitative perspective on the mostly unexplored phenomena of adsorption of FN on HA in presence of an external stimulus.

## 2. Simulation details

The structure of the model protein, FN-III_10_ was taken from PDB database (code: 1TTF). FN-III_10_ consists of 94 residues with a molecular weight of 9.92 kDa. It has net zero charge, together with total dipole moment of 245.7 D. This module is of particular interest, because of the presence of an integrin binding site (RGD loop) in it.^15^ The rectangular HA slab structure was obtained from the database of INTERFACE force field (https://bionanostructures.com/interface-md/). This rectangular unit cell of HA features the following lattice parameters: *a=*9.4170 Å, *b=* 16.3107 Å, *c=*6.8750 Å, α=β=γ=90°.

A HA slab of 7.5336×9.7864×4.7500 nm^3^ was built and used as a biomaterials substrate in the MD simulation. A periodic box of size 7.5336×9.7864×14.04 nm^3^ was solvated with TIP3P water. The system contains 25,965 water molecules and an overall 96715 atoms. Initially, the protein molecule was placed 5 Å above the HA (001) facet and the preferred adsorption orientation of the protein was chosen from the previous report.^16^ Energy minimization of the whole system was carried out in 10,000 steps by means of the steepest descent algorithm to remove close contacts between molecules. Then, the system was heated to 300 K in 1 ns. Modified Berendsen thermostat^24^ was employed for temperature coupling (at 300 K) with a coupling constant of 0.1 ps. Additional 2 ns equilibration MD was carried out in NPT ensemble with a reference pressure of 1.0 bar. Parrinello-Rahman barostat^25^ was engaged for pressure coupling with coupling constant of 0.5 ps. Position restraints were applied on both protein and HA slab during both equilibration stages in NVT and NPT ensembles. The coupling time of barostat was increased to 2.0 ps during production runs. 2 fs was chosen as the integration time step throughout the simulation. External electric fields of upto 1.00 V/nm were applied along parallel and anti-parallel to z-axis. Throughout this article, positive and negative electric field corresponds to electric field applied parallelly and anti-parallelly to z-direction, respectively. Position restraints on HA surface was only kept intact during production run. Data from 100 ns MD runs were collected for different kind of analysis. All MD simulations were carried out in GROMACS 5.1.4 software,^26^ using AMBER99SB ILDN force field.^27^ Force field parameters for HA was taken from INTERFACE force field.^28^ Necessary changes (Kcal/mol to kJ/mol and Å to nm; σ_GROMACS_ =(σ_INTERFACE_)/2^1/6^) were done to make INTERFACE force field parameters compatible with GROMACS. LINCS algorithm^29^ was employed for bond constraints (all bonds). Particle Mesh Ewald (PME) was used to calculate the long-range electrostatic interaction.^30^ A cut-off distance of 1.0 nm was adopted for short-range non-bonded interactions. The calculations of interaction energy, root mean square deviation (RMSD), dipole moment, hydrogen bonds, protein surface distance were carried out with the help of different GROMACS modules. VMD 1.9.3 software^31^ was used for visualization and salt bridge analysis. The cut-off used for salt bridge and hydrogen bond analysis was set to 3.2 Å and 3.5 Å, respectively. The percentage of occurrence of salt bridges was calculated from the relation N_on_/N_total_, where N_on_ is the number of frames where the salt bridge was formed and N_total_ was the total number of frames.^32^

## 3. Results

We have employed all-atom MD simulation to probe into FN-HA interactions in reference to temporal evolution of interaction energy, dipole moment, hydrogen bonds and other response parameters of the simulated system (Fig. S1 in supplementary information). In the context of the cell-material interaction, the position of the RGD loop and its accessibility for integrin protein is of particular interest. As iterated before, integrin, one of the major cell surface receptor protein, interacts with fibronectin through specific amino acid sequence and this interaction leads to the formation of focal adhesion complex (Fig. S1 in supplementary information).^15^ In this section, the adsorption behavior of FN is described in reference to the simulation results. The biophysical significance of the results will be described in the ‘Discussion’ section.

### 3.1 FN-HA interaction energy

At the molecular level, the FN-HA interaction can be assessed by the temporal evolution of the distance of separation between center of mass (COM) of FN module and that of HA slab. This average distance is a signature of the occurrence of protein adsorption on the material surface. External electric field dependent COM-COM average distance has been plotted in Fig. 1. For positive and low field strength (upto 0.50 V/nm), the average distance remains within the range of 3.1 nm to 3.3 nm, throughout the simulation time, indicating adsorbed state of protein (Fig 1). When the field was not applied, distance between protein and surface slightly increased from 3.1 nm at ∼3 ns upto 3.4 nm at ∼27 ns. Oscillatory nature was recorded after this time upto ∼73 ns and a steady adsorbed state was obtained for the remainder of the simulation duration (Fig. 1). In case of 0.25 V/nm, the protein got adsorbed on the HA surface around 20 ns. A bump arose near 80 ns and the protein surface distance further got reduced to 3.1 nm (Fig. 1). At increased field strength (0.50 V/nm), a nearly constant protein surface distance of nm was maintained from ∼54 ns. For 0.75 V/nm, although the initial distance was around 3.35 nm upto ∼40 ns of simulation, it started to increase after that. The distance reached 3.45 nm at ∼63 ns. The increment of COM-COM distance implies a comparatively weak adsorbed state. At 1.00 V/nm, the protein moved away from the surface within first 8 ns of simulation and maintained a nearly constant distance of ∼4.2 nm for the rest of the simulation time. In this case, the protein is loosely bound to the surface.

**Fig. 1:**
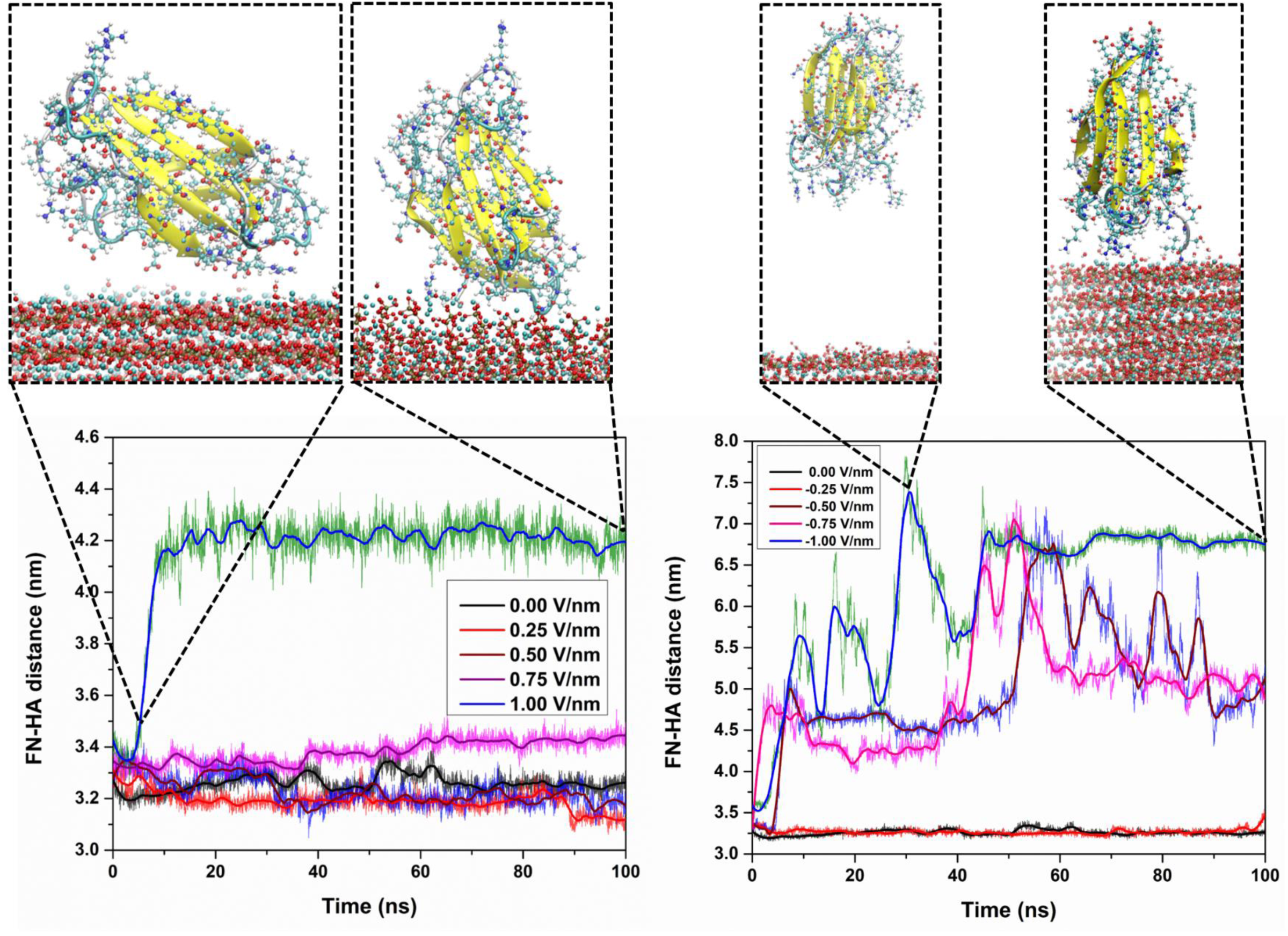
Electric field application and its directionality influence the adsorption kinetics. MD results to illustrate the temporal evolution of distance between centre of mass (COM) of fibronectin and that of hydroxyapatite (HA) substrate. The larger distance indicates that, high field strength does not favour FN adsorption on HA surface. In the graphs, the legend colour codes represent the thick lines (generated using Savitzky–Golay filter). Snapshots of trajectory at different time points have been shown for ±1.00 V/nm. Water molecules are not shown due to clarity. The secondary structure of FN has been represented with colour code: Yellow: β sheet, pale blue: turns, white: other residues. FN and HA have been presented using CPK colouring scheme with colour code: Red: O, pale blue: C(FN) and Ca (HA), white: H, blue: N, golden yellow: P.

When field was applied anti-parallel to the z-direction (negative z-direction), FN remained adsorbed on the surface for an applied field strength upto -0.25 V/nm (Fig. 1). At this field strength, protein surface distance was almost constant (∼3.25 nm) throughout the simulation time. With further increase of field strength, the COM of protein moved away from the surface and the adsorbed state became weaker (Fig. 1). At field strength beyond -0.25 V/nm, FN, at some time points, completely desorbed from the surface and again resorbed. The jumps in the distance profile at high electric field represent the desorbed state of FN (Fig. 1). For a field strength of -1.00 V/nm, the desorbed state of FN remained for first ∼40 ns of the simulation. The different behavior of the COM-COM distance at different field strength indicates towards the influence of external electric field on the adsorption phenomena of protein on materials surface.

It is equally important to understand the modes of protein-HA interaction. FN-HA interaction energies at different electric field strengths have been calculated using GROMACS module and are represented in Fig. 2. It is evident that the long range attractive electrostatic interaction dominates fibronectin adsorption on the HA surface. In the absence of external field, the average electrostatic interaction between protein and HA was around -426.2±44.8 kJ/mol for the first 21ns. An increment was recorded afterwards upto ∼54 ns, followed by a sharp fall. The interaction energy remained constant at ∼ -628.8±38.2 kJ/mol from 80 ns onwards. When the field strength was increased to 0.25 V/nm, a nearly steady state with an average interaction energy value of ∼549.4±44.3 kJ/mol was reached at ∼50 ns and a further decrease was noted after 89 ns. A slight decrease in the interaction energy was observed at an electric field strength of 0.5 V/nm, whereas a nearly constant energy value of ∼ -403.6±53.1 kJ/mol was recorded from ∼60 ns at field strength of 0.75 V/nm. Coulomb interaction energy between FN and HA fluctuates with a mean value of ∼ -245.2±5.8 kJ/mol for 1.00 V/nm.

**Fig. 2:**
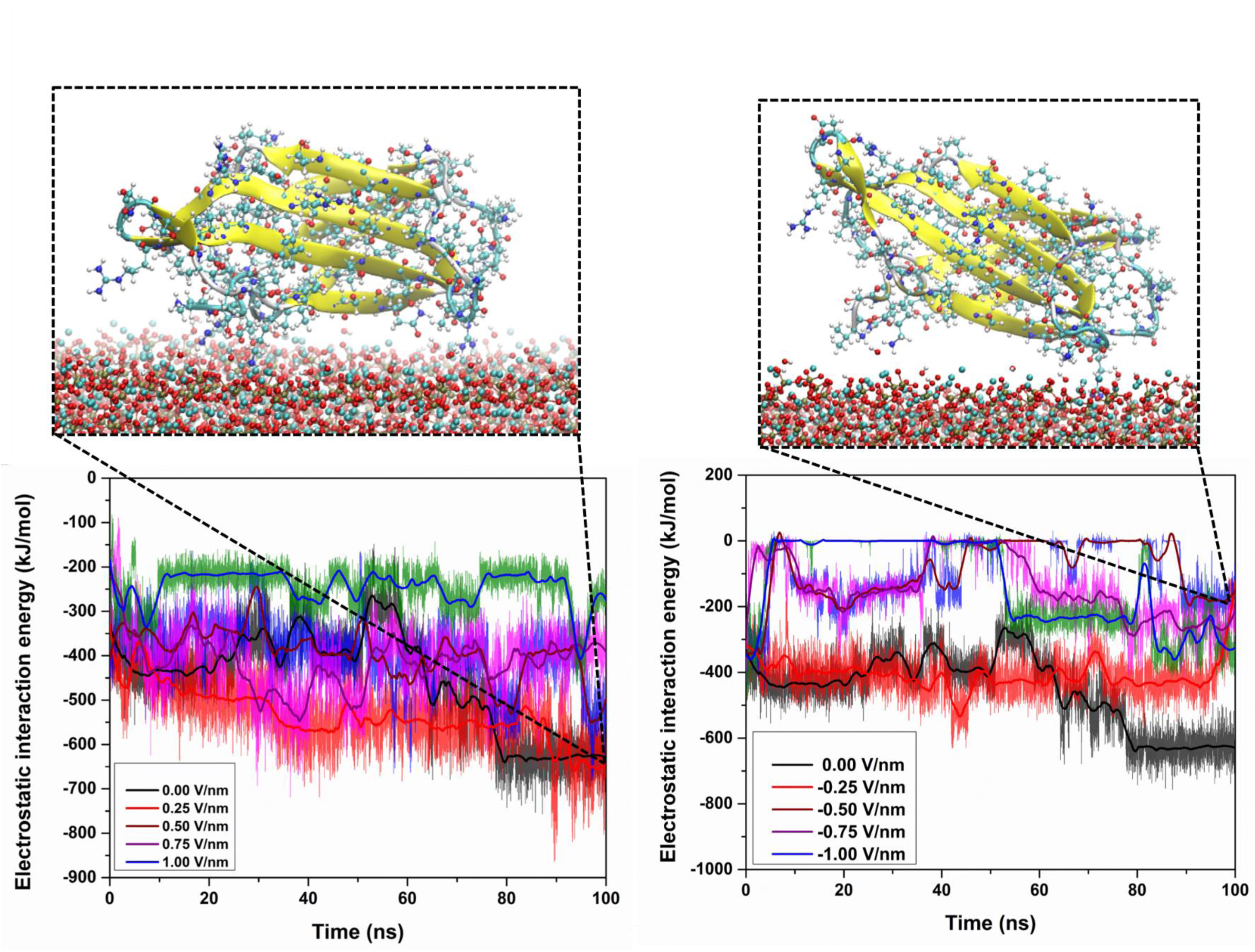
FN-HA interaction is dominated by the long-range coulombic interaction. Time evolution of the electrostatic interaction energy between the FN-HA for different electric field strength. Interaction strength generally reduces at high field strength, which represents weakly adsorbed state of FN. In the graphs, the legend colour codes represent the thick lines (generated using Savitzky–Golay filter). Final configurations (at 100 ns) have been shown for ±0.25 V/nm. Water molecules are not shown due to clarity. The secondary structure of FN has been represented with colour code: Yellow: β sheet, pale blue: turns, white: other residues. FN and HA have also been presented using CPK colouring scheme with colour code: Red: O, pale blue: C(FN) and Ca (HA), white: H, blue: N, golden yellow: P.

When field direction is reversed, the situation was quite similar, only upto -0.25 V/nm. The protein remained adsorbed on the surface throughout the simulation time (steady interaction energy). At higher field, the interaction energy decreased further and sometimes approached to zero during the investigated time window (Fig. 2). These low interaction energies at high field correspond to a desorbed state of the protein.

The time averaged electrostatic interaction energy has been listed in Table 1. With the application of positive electric field, the average interaction energy first increased compared to that in the absence of external field (Table 1). Beyond +0.25 V/nm, it decreased non-monotonously. On the other hand, the interaction energy started to decrease from the beginning, when the field direction is reversed. It increased at -0.75 V/nm and -1.00 V/nm. Even at the same magnitude of the field strength, the interaction energy was noted to be less, when the field is applied anti-parallel to z-direction, except at -1.00 V/nm. The change in the attractive interaction energy establishes the fact that, although FN adsorption was promoted at low field strength (up to 0.25 V/nm), application of high electric field does not favor protein adsorption on HA surface.

**Table 1:** Average interaction energy between FN and HA, as calculated from the trajectory obtained from MD simulation.

| Electric field strength (V/nm) | Coulomb energy (kJ/mol) | Error (kJ/mol) |
| --- | --- | --- |
| **-1.00** | -363.2 | 150 |
| **-0.75** | -139.1 | 38.0 |
| **-0.50** | -92.3 | 30.0 |
| **-0.25** | -408.2 | 9.8 |
| **0.00** | -454.3 | 50.0 |
| **0.25** | -533.0 | 25.0 |
| **0.50** | -392.3 | 19.0 |
| **0.75** | -413.2 | 18.0 |
| **1.00** | -245.2 | 5.8 |

Another important mode of interaction is mediated via the formation of hydrogen bonds between the material surface and fibronectin, which has been penned in section S1 of the supplementary section.

For different electric field strength, the number of contacts between HA and FN has been presented in Fig. 3, for last 10 ns of the total simulation time. The atom contact number was observed to reach the highest at +0.25 V/nm field strength (Fig. 3). For higher +ve field strength, the number of contacts decreased, and the lowest values were recorded for +1.00 V/nm. When field direction was reversed, the number of contacts was found to be lesser for all field strength, compared to that in absence of any external field (Fig. 3). In corroboration with previously obtained results, the reduced number of H-bonds and number of contacts at very strong field intensity represents the weakly adsorbed state of FN on HA.

**Fig. 3:**
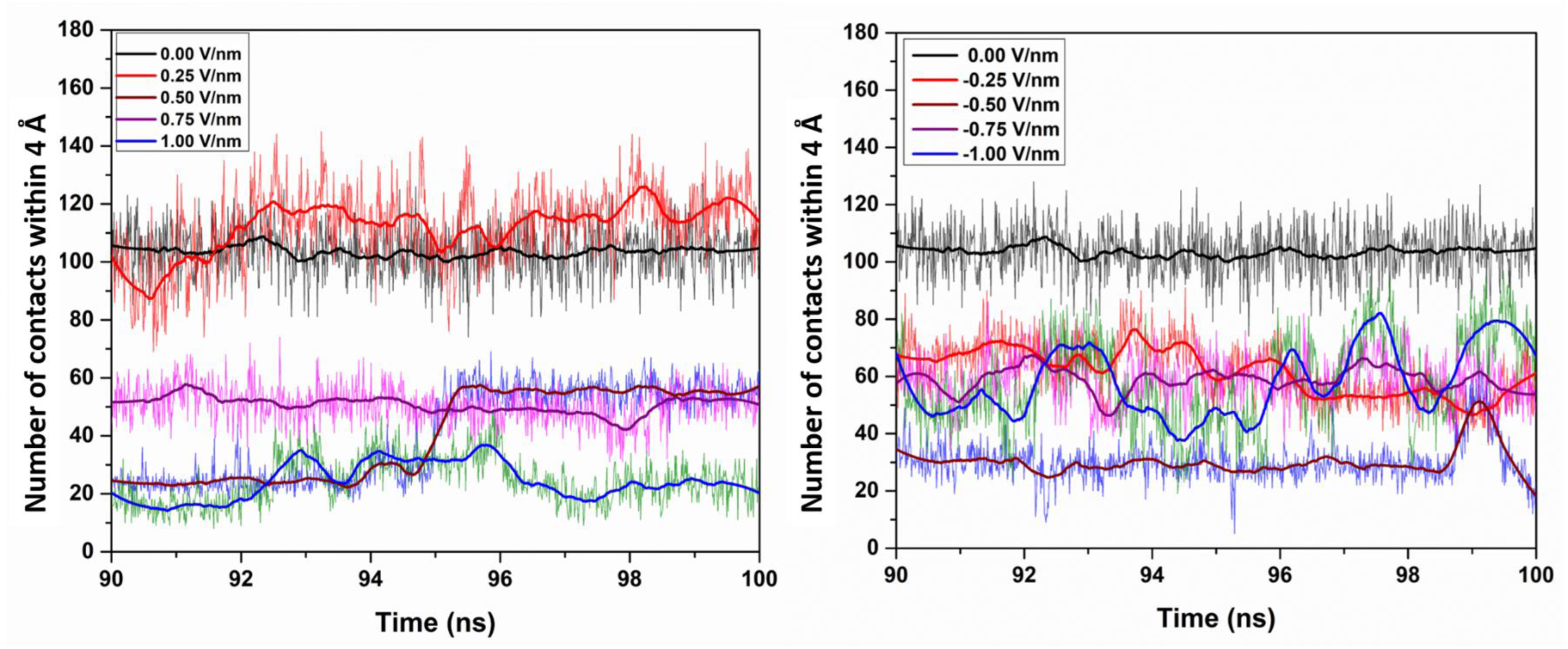
Larger number of contacts is an indication of adsorption and better indication. Time evolution of number of contacts between FN and HA at different electric field strength and direction. Highest number of contacts was formed in case of +0.25 V/nm, which can be correlated with the observed highest interaction strength between FN and HA at this field strength. A contact is defined when two atoms belonging to FN and HA are within a distance of 4Å. In the graphs, the legend colour codes represent the thick lines (generated using Savitzky–Golay filter).

### 3.2 Probing into absorbing residues and secondary structure

It is a well-known fact that, in the course of the adsorption process, proteins change their native structure, depending on the nature of the interaction with an inorganic surface.^1-3^ As mentioned before, the conformational change of adsorbed protein is important, because it affects cell-material interaction. Only a few types of residues of any protein interact with the inorganic surface, depending on the nature of surface bonds and the amino acids. The interacting residues of FN, at different electric field strengths, together with their closest groups, are listed in Table Arginine, aspartic acid, glycine, glutamic acid, etc. are the dominant residues that are found near the HA surface, in agreement with the existing literature.^16^ Among them, positively charged residues like Arginine and Lysine interact via NH_2_/NH_3_^+^ groups. Negatively charged residues (glutamic acid and aspartic acid) and neutral amino acids (glycine) were observed to interact with the HA surface through COO^-^ groups. A notable observation is that, beyond field strength of 0.25 V/nm, the number of interacting residues declined. Moreover, Arg93 residue ceased to interact with HA surface at higher electric field strength. This effect was prominent when the field was applied anti-parallel to z-axis. In a previous report, this residue (Arg93) has been noted as the most strongly interacting residue with HA.^16^ Therefore, withdrawal of Arg93 from HA-FN interaction at higher field strengths reflects weaker adsorbed state. Another notable observation is that the number of interacting residues declined when the electric field was applied along –ve z direction (Table 2). The change in the adsorbing residues with the change in field strength and direction is a signature of the profound effects of external perturbation like electric field on adsorption kinetics.

**Table 2:** Summary of various residues of FN adsorbed on HA surface, in presence of electric field.

| Electric Field strength (V/nm) | Anchoring residues (nearest group) |
| --- | --- |
| -1.00 | Val1(NH<sub>3</sub><sup>+</sup>), Ser2 |
| -0.75 | Arg78 (NH<sub>2</sub>) |
| -0.50 | Arg78 (NH<sub>2</sub>) |
| -0.25 | Asn42 (NH<sub>2</sub>), Gly40 (COO<sup>-</sup>) Asp67 (COO<sup>-</sup>), Arg93 (NH<sub>2</sub>), |
| 0.00 | Val1 (NH<sub>3</sub><sup>+</sup>), Glu9(COO<sup>-</sup>), Asp7 (COO<sup>-</sup>), Arg93 (NH<sub>2</sub>), |
| 0.25 | Arg6 (NH<sub>2</sub>), Asp3 (COO<sup>-</sup>), Asp7 (COO<sup>-</sup>), Lys86 (NH<sub>3</sub><sup>+</sup>), Arg93 (NH<sub>2</sub>) |
| 0.50 | Asp7 (COO<sup>-</sup>), Glu9 (COO<sup>-</sup>), Arg93 (NH<sub>2</sub>) |
| 0.75 | Arg93(CH<sub>2</sub>), Glu9(COO<sup>-</sup>), Val11 (COO<sup>-</sup>), |
| 1.00 | Gly65 (α-C) Pro64 (part of side chain) |

The relevance of RGD loop in the context of cell-material interaction has been mentioned before (Fig.S1 in supplementary). The position of the RGD loop at different field strengths and directions has been shown in Fig. 4. In the absence of an electric field, the RGD loop has been exposed in the solution and is available for cell-material interaction (Fig. 4). When E-field was applied parallel to +z-direction, RGD loop remained exposed to the solvent for all field strength (Fig. 4). Upto -0.25 V/nm, RGD sequence was away from the HA surface (Fig. 4). At field strength of -0.50 and -0.75 V/nm, one of the residues (Arg78) of the RGD loop was adsorbed on the surface. At -1.00 V/nm, the RGD loop remained very close to the surface (Fig. 4). These configurations, therefore, will not promote cell material interaction. Therefore, it can be commented that, cell-material interaction can be modulated by changing electric field strength and direction.

**Fig. 4:**
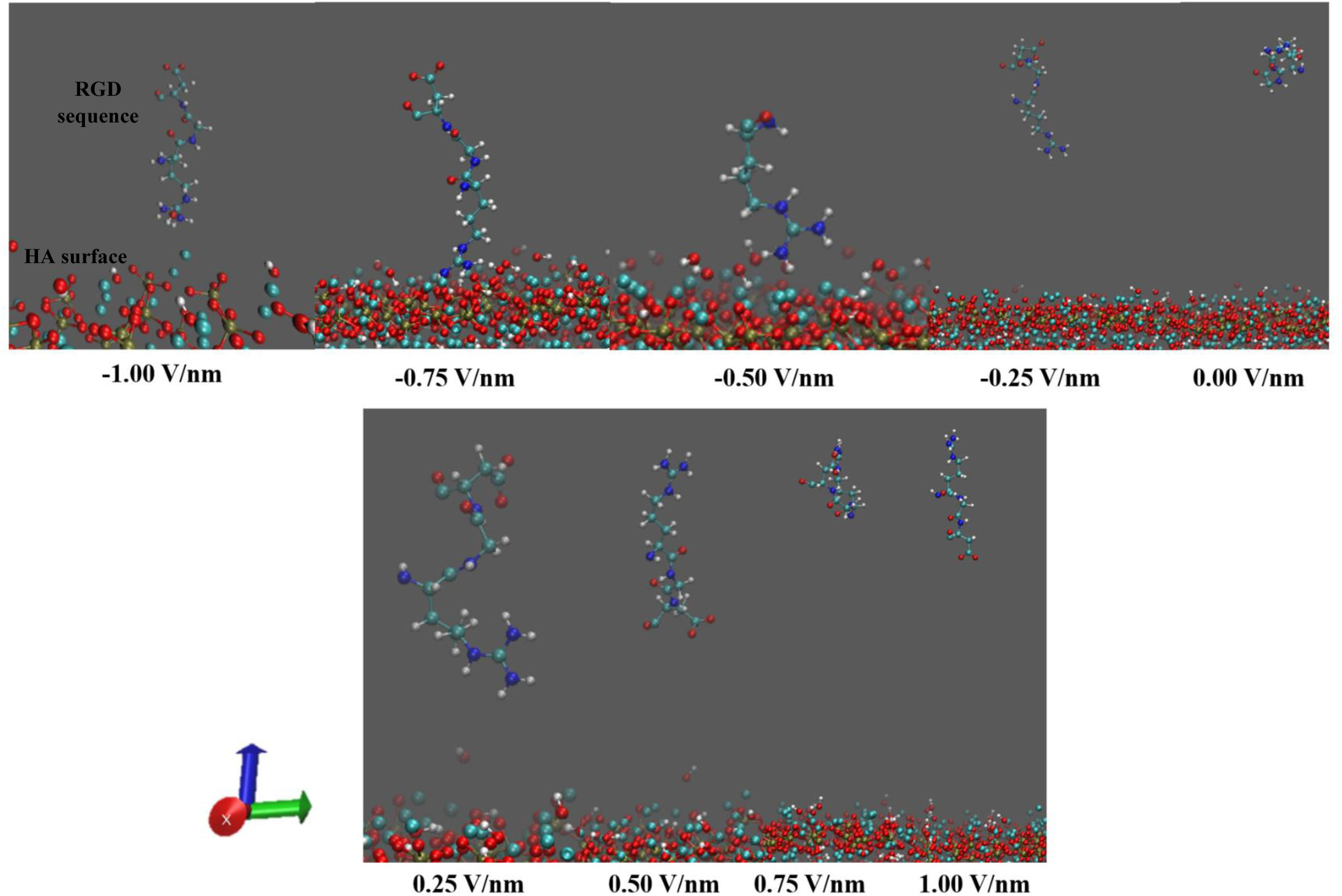
Accessibility of RGD loop profoundly influences cell-material interaction. It is noteworthy that, conformation with RGD loop exposed to the solvent is suitable for cell-material interaction. Position of RGD loop w.r.t. HA surface after 100 ns of simulation at different electric field. RGD loop is not exposed to solvent at negative field direction. Hence, application of negative field is likely to hinder cell adhesion on HA surface (Atomic colour code: Red: O, blue: N, pale blue: C (FN) and Ca (HA), white: H, golden yellow: P. Axis colour code: Red: x, green: y, blue: z).

In order to get a qualitative idea about the conformal changes, the aligned initial and final structure have been presented in Fig. S4 in supplementary information. It can easily be seen that the structural changes took place mainly in the flexible loop region of the protein. Not much significant changes were noticed in the β-sheet structure, except at -1.00 V/nm (Fig. S4 in supplementary information), because of its unique structural feature. There exists a distinguished pattern of inter-residual H-bonds in the β-sheet, which provides a high structural stability. Hence, it is very difficult to alter this structure by means of any external stimuli.

In order to obtain a quantitative estimate of the structural changes, the percentage of the β-sheets was measured, and the number lies between 36 to 58% (Fig. 5). For positive field direction, percentage of β-sheets increased upto 0.75 V/nm and then it decreased. For negative direction, the increasing trend was observed upto -0.50 V/nm (Fig. 5). Additionally, RMSD of the backbone of the protein was calculated and is shown in Fig. 6. It is noteworthy that, the RMSD first decreased with the application of electric field (+0.25 V/nm) and beyond 0.25 V/nm, it started to increase, irrespective of the field direction (Fig. 6). Besides this, at high field strength, RMSD was observed to be higher, when the field was applied anti-parallel. This indicates towards the dependency of conformal change of FN not only on electric field strength, but also on its direction.

**Fig. 5:**
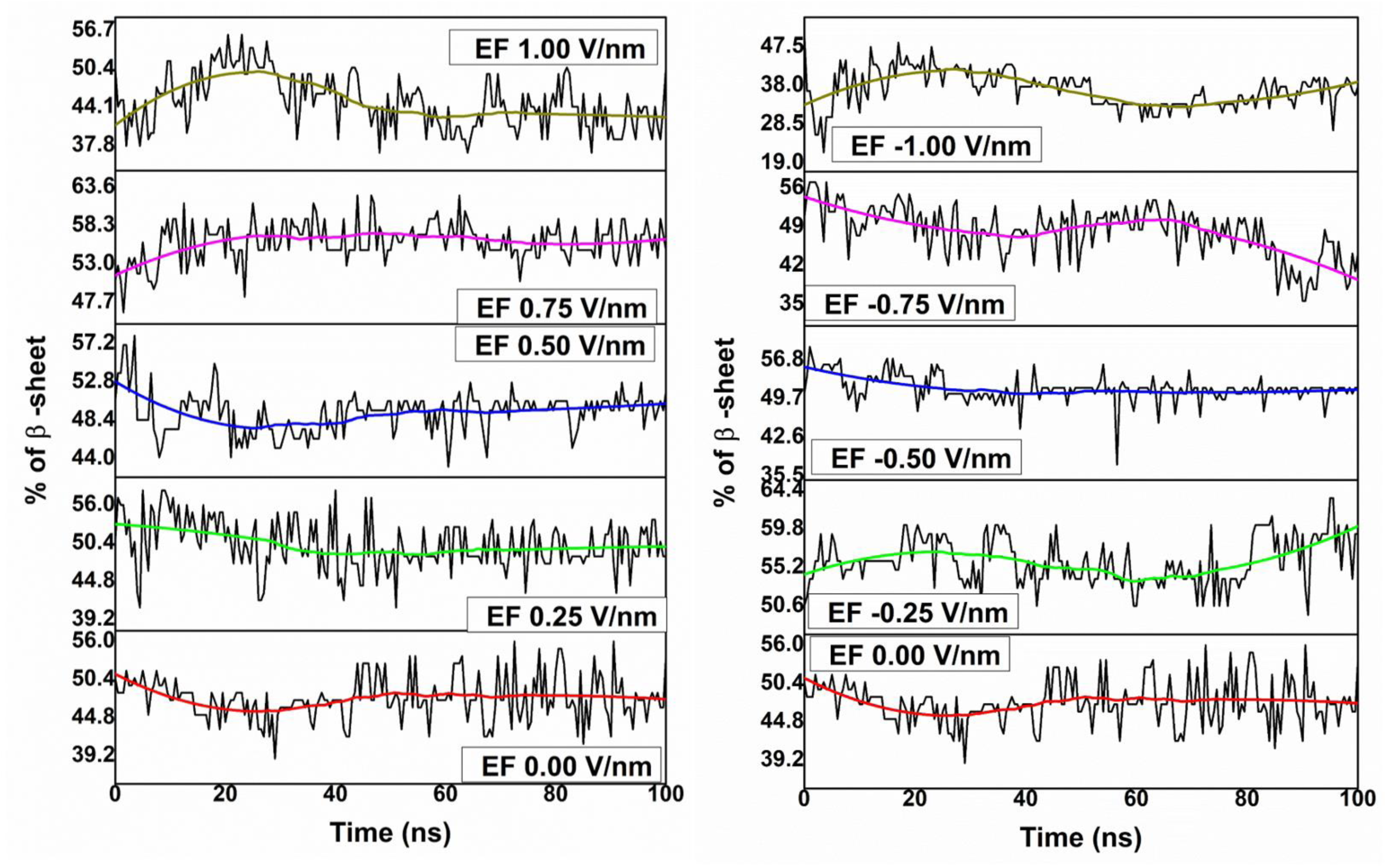
Electric field application affects the structural integrity of fibronectin. Percentage of β-sheet as a function of simulation time, at different electric field strength and direction. The stable β-sheet structure was not affected significantly at low field strength and positive direction. Therefore, FN structure cannot be greatly altered by using low electric field intensity. Notable change occurred at -1.00 V/nm. Smoothed thick lines were generated using Savitzky– Golay filter.

**Fig. 6:**
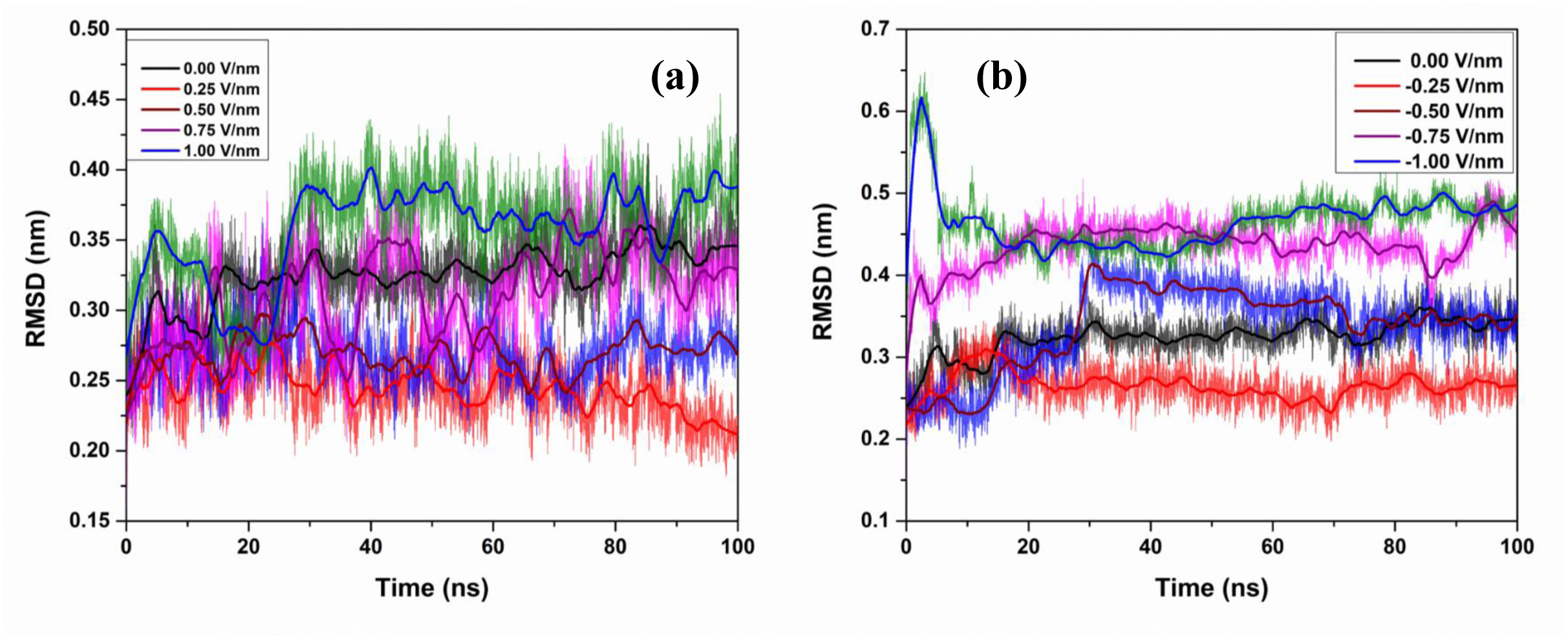
Structure of FN significantly deviates from initial structure in presence of electric field. Root mean square deviation (RMSD) of the FN-backbone as a function of (a) positive and (b) negative electric field strength. For both +ve and -ve field strength, RMSD first decreases, and then increases with the magnitude of electric field. The change in RMSD points out towards the fact that, the electric field does not affect the FN structure in a drastic manner. Maximum changes occur in the flexible loop areas (see article for details). In the graphs, the legend colour codes represent the thick lines (generated using Savitzky–Golay filter).

The shape of protein can be approximated by an ellipsoidal shape, defined by the principal moments of inertia and principal axes.^33^ The principal moments of inertia of FN and their ratios has been listed in Table 3. From the table, it can be easily seen that, when electric field was applied parallel to the z-axis, I_1_ and I_2_ increased upto 0.50 V/nm and then decreased. An opposite trend was noted for I_3._ The ratio, I_1_/I_3_ also exhibited the similar pattern, I_1_/ I_2_. When field direction was reversed, the increasing trend was noticed only upto -0.25 V/nm for I_1_, I_2_ and I_1_/I_3._ As before, I_3_ exhibited opposite trend. Various parameters calculated from the simulated trajectories to probe the structural change of FN clearly demonstrated that, the electric field did not profoundly affect the structural integrity of the protein, probably due the presence of stable β-sheet structure, as mentioned before.

**Table 3:** Principal components of moment of inertia of FN and their ratios

| **Electric field(V/nm)** | **I<sub>1</sub> (amu Å<sup>2</sup>)** | **I<sub>2</sub> (amu Å<sup>2</sup>)** | **I<sub>3</sub> (amu Å<sup>2</sup>)** | **I<sub>1</sub>/I<sub>2</sub>** | **I<sub>1</sub>/I<sub>3</sub>** | **I<sub>2</sub>/I<sub>3</sub>** |
| --- | --- | --- | --- | --- | --- | --- |
| -1 | 2441743.5 | 2323944.25 | 1515877.125 | 1.050689 | 1.610779 | 1.533069 |
| -0.75 | 2192289 | 2115115.75 | 1412201.25 | 1.036487 | 1.552391 | 1.497744 |
| -0.5 | 2393993.25 | 2299658 | 1109157.25 | 1.041021 | 2.158389 | 2.073338 |
| -0.25 | 2605389.75 | 2515012.5 | 1071420.25 | 1.035935 | 2.431716 | 2.347363 |
| 0 | 2439327.25 | 2366321.75 | 1248351 | 1.030852 | 1.95404 | 1.895558 |
| 0.25 | 2670087.5 | 2489611.5 | 1078987 | 1.072492 | 2.474624 | 2.30736 |
| 0.5 | 2703543.5 | 2608899 | 1036026.063 | 1.036278 | 2.609532 | 2.518179 |
| 0.75 | 2519970 | 2372511.5 | 1142999.5 | 1.062153 | 2.204699 | 2.075689 |
| 1 | 2352267.75 | 2191058 | 1196919.625 | 1.073576 | 1.965268 | 1.830581 |
| no surface | 1696709 | 1588183.375 | 699086 | 1.068333 | 2.427039 | 2.2718 |

One of the ways to determine the structural integrity of proteins is to analyze the intramolecular salt bridge network, as it contributes significantly towards the thermostability of proteins.^34^ The number of salt bridges at different electric field and the corresponding residues are presented in Table 4. When no external field was applied, Arg6 formed salt bridges with Asp3, Asp7 and Asp23, resulting in a salt bridge network. Another salt bridge network observed was Arg6-Asp3-Lys86 (Table 4). These two salt bridge nets together with other salt bridges are retained at field strength of +0.25 V/nm. Moreover, a new salt bridge was observed between Asp80 and Arg30, leading to a new salt bridge net consisting of Arg78-Asp80-Arg30. These networks were affected with the increment of the field strength. At 0.50 V/nm, a new network, Asp80-Arg33-Glu47 was observed and the salt bridge between Arg78-Asp80 and Asp3-Lys86 was broken. More salt bridges were formed with the application of stronger electric field (Table 4). The presence of Asp3-Lys86-Arg6 and Asp23-Arg6-Asp3 salt bridge networks was recorded in both 0.75 V/nm and 1.00 V/nm. Additional formation of salt bridges among Asp80-Arg33-Glu47 and Glu9-Arg93-Asp67 were observed at 1.00 V/nm.

**Table 4:** Salt bridge forming residues of FN at different electric field strength.

| Electric field strength (V/nm) | Salt bridge forming residues |
| --- | --- |
| -1.00 | Asp23-Arg6<br>Asp7-Arg93<br>Asp3-Lys86<br>Glu9-Arg93<br>Asp7-Lys54<br>Asp7-Arg6<br>Asp23-Lys54<br>Glu47-Arg33<br>Asp3-Arg6<br>Asp80-Arg33 |
| -0.75 | Asp80-Arg33<br>Asp23-Arg6<br>Asp7-Arg93<br>Glu9-Arg93<br>Asp7-Arg6<br>Asp23-Arg93<br>Asp3-Arg6<br>Glu47-Arg33<br>Asp3-Lys86<br>Glu9-Lys63<br>Asp23-Lys54<br>Asp7-Lys63 |
| -0.50 | Asp23-Arg6<br>Glu47-Arg33<br>Asp3-Lys86<br>Asp7-Arg6<br>Asp80-Arg30<br>Asp80-Arg78<br>Asp3-Arg6<br>Glu38-Lys63<br>Asp67-Arg93<br>Asp3-Arg78<br>Asp80-Arg33 |
| -0.25 | Asp23-Arg6<br>Glu47-Arg33<br>Asp3-Lys86<br>Asp67-Arg93<br>Asp7-Arg6<br>Asp80-Arg30<br>Glu38-Lys63<br>Asp3-Arg6 |
| 0.00 | Asp23-Arg6<br>Asp80-Arg78<br>Asp3-Lys86<br>Glu47-Arg33<br>Asp67-Arg93 |
|  | Asp7-Arg6<br>Glu38-Lys63<br>Asp3-Arg6 |
| 0.25 | Asp80-Arg78<br>Glu47-Arg33<br>Asp3-Lys86<br>Asp80-Arg30<br>Asp7-Arg6<br>Glu38-Lys63<br>Asp67-Arg93<br>Asp3-Arg6<br>Asp23-Arg6 |
| 0.50 | Asp80-Arg33<br>Glu47-Arg33<br>Asp80-Arg30<br>Asp67-Arg93<br>Asp23-Arg6<br>Glu38-Lys63<br>Asp3-Arg6 |
| 0.75 | Asp80-Arg78<br>Asp23-Arg6<br>Asp3-Lys86<br>Glu47-Arg33<br>Asp67-Arg93<br>Asp7-Arg6<br>Glu38-Lys63<br>Asp3-Arg6 |
| 1.00 | Asp80-Arg33<br>Asp3-Lys86<br>Glu47-Arg33<br>Glu9-Arg93<br>Asp67-Arg93<br>Asp80-Lys86<br>Asp23-Arg6<br>Glu38-Lys63<br>Asp3-Arg6 |

A noteworthy observation is that more salt bridges were formed in the presence of strong electric field, when field direction is reversed (Table 4). Salt bridge networks, formed in the presence of negative electric fields, were also noticed to be marginally different from those appeared in the presence of the positive electric field. For example, Lys54 residue formed salt bridge with Asp23 at -0.75 V/nm and with both Asp23 and Asp7 at -1.00 V/nm. This particular residue did not take part in salt bridge formation in presence of positive electric field. Asp7-Arg93-Glu9 is another such salt bridge net, which was formed only in association with an anti-parallel electric field. The temporal evolution of these salt bridges in an interesting facet of study to analyse the stability of FN structure. The details of this particular aspect have been included in section S2 of the supplementary section.

### 3.3 Dipole moment distribution

Dipole moment of the various system components plays an important role in the adsorption process. The dipole moment orientation of fibronectin has been shown in Fig. 7(a)-7(b). The total dipole moment of protein after 100 ns of simulation and the corresponding angle with the z-axis, has been listed in Table 5. From Table 5, it is clear that the total dipole moment of the protein increased with an increase in field strength, irrespective of the direction. Also, the dipoles tended to align themselves with the field direction. The z-component of the total dipole moment of fibronectin has been plotted in Fig. 7(c)-7(d). From the graphs, it can easily be noticed that, although the magnitude of z-component of the dipole moment increased with the increment of the field strength, the increment depended on the field direction (Fig 7(c)-7(d)). An increment of the dipole moment took place within 100 ps of the simulation time (Fig. 7). The other components (x and y) of dipole moment vector of FN did not exhibit much significant change (Fig. S8 in supplementary section).

**Table 5:** Dipole moment of FN at different electric fields after 100 ns of simulation and their orientation relative to z-axis.

| Electric field strength (V/nm) | Dipole moment (debye) | $\cos\theta$ | Angle (Degree) |
| --- | --- | --- | --- |
| -1.00 | 751.9 | -1.00 | 180.0 |
| -0.75 | 689.7 | -1.00 | 180.0 |
| -0.50 | 500.1 | -0.99 | 171.9 |
| -0.25 | 233.2 | -0.05 | 92.9 |
| 0.00 | 225.8 | 0.22 | 77.3 |
| 0.25 | 228.6 | 0.32 | 71.3 |
| 0.50 | 302.4 | 0.83 | 33.9 |
| 0.75 | 418.1 | 0.91 | 24.5 |
| 1.00 | 589.0 | 1.00 | 0.0 |

**Fig. 7:**
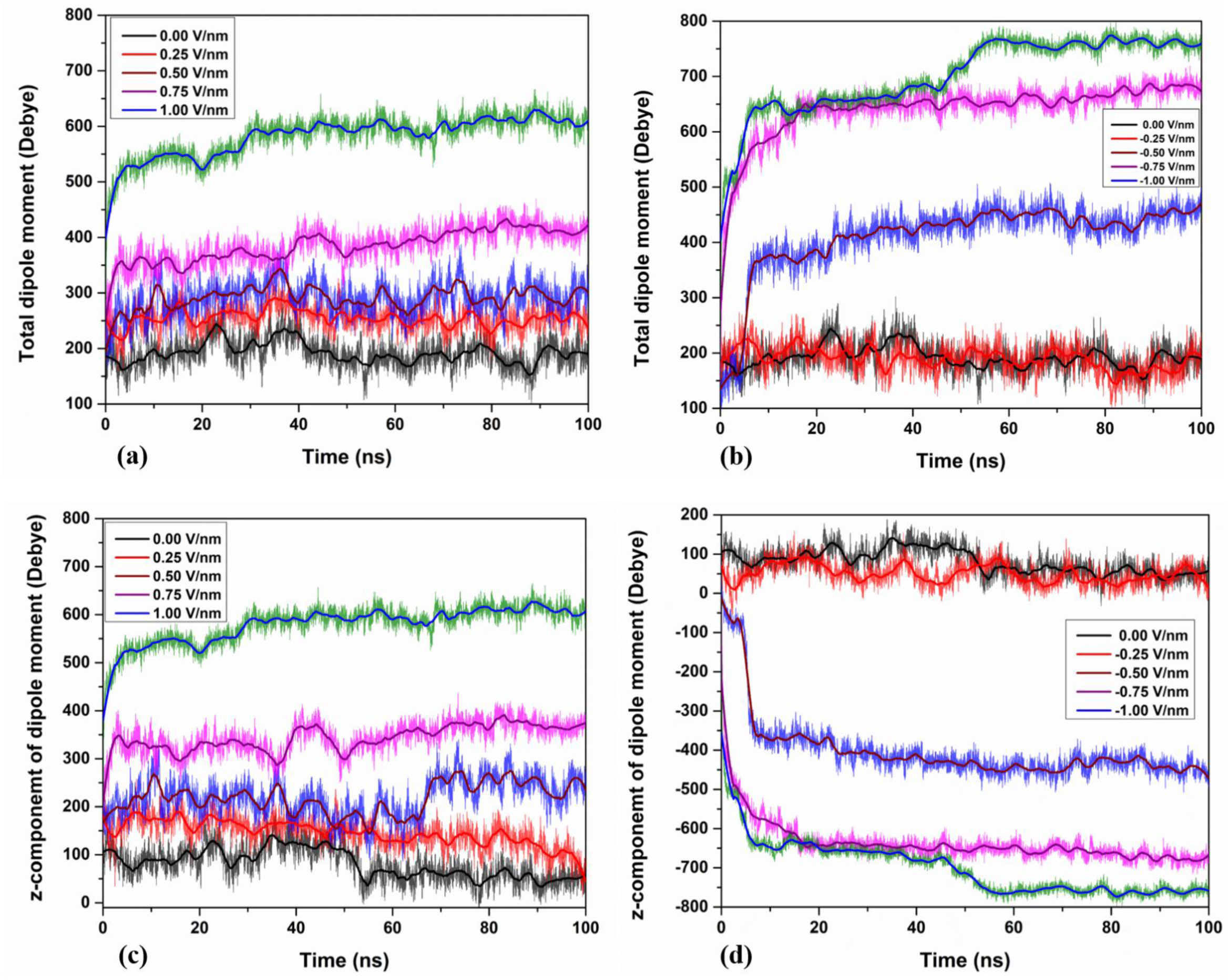
Dipole moment of protein changes with magnitude and direction of external electric field. Time evolution of (a)-(b) total and (c)-(d) z-component of dipole moment of protein at different electric fields. The change in total dipole moment with the increase of field strength can be corroborated with the rearrangement of local charges of FN. The recorded trend of the z-component of dipole moment indicates that, the FN dipole moment vector is orientating itself along the field direction. In the graphs, the legend colour codes represent the thick lines (generated using Savitzky–Golay filter).

Total dipole moment of HA after 100 ns simulation time has been tabulated in Table 6. When the field was applied along the z-direction, magnitude of the total dipole moment of HA decreased first and then it began to increase (Table 6). On the other hand, a monotonous increasing trend was noted, when the field direction was reversed. Moreover, like fibronectin, the angle between dipole moment and z-axis decreased and increased monotonically with positive and negative electric fields, respectively (upto 0.75 V/nm). This trend implies that the dipole moment vector of HA was trying to align itself with the field direction, just like the protein. The z-component of dipole moment also followed similar pattern (Fig. S9 in supplementary information).

**Table 6:** Dipole moment of bulk HA surface as a function of electric field strength and direction at time point of 100 ns. Angle made by the dipole moment vector with z-direction has also been shown.

| Electric field strength (V/nm) | Dipole moment (debye) | $\cos\theta$ | Angle (Degree) |
| --- | --- | --- | --- |
| -1.00 | 17948.4 | -0.82 | 145.1 |
| -0.75 | 13847.4 | -0.92 | 156.9 |
| -0.50 | 12491.2 | -0.71 | 135.2 |
| -0.25 | 7947.8 | -0.64 | 129.8 |
| 0.00 | 8180.6 | -0.35 | 110.5 |
| 0.25 | 8191.3 | -0.08 | 94.6 |
| 0.50 | 8448.4 | 0.27 | 74.3 |
| 0.75 | 11123.2 | 0.66 | 48.7 |
| 1.00 | 18954.2 | 0.64 | 50.2 |

The electrical behavior of water was found to follow a similar trend, as depicted in Table 7. In this case, the angle between the dipole moment vector and the z-axis decreased with the increment of the field strength, when applied parallel to the z-direction (Table 7). On the other hand, the very same angle remained constant, when the field direction is reversed. The magnitude of the dipole moment vector increased with the field strength, irrespective of the direction, except for -0.25 V/nm (Table 7). The z-component of dipole moment exhibited similar trend like FN/HA. It increased with the positive electric field and decreased with the negative field (Fig. S10 in supplementary information).

**Table 7:**
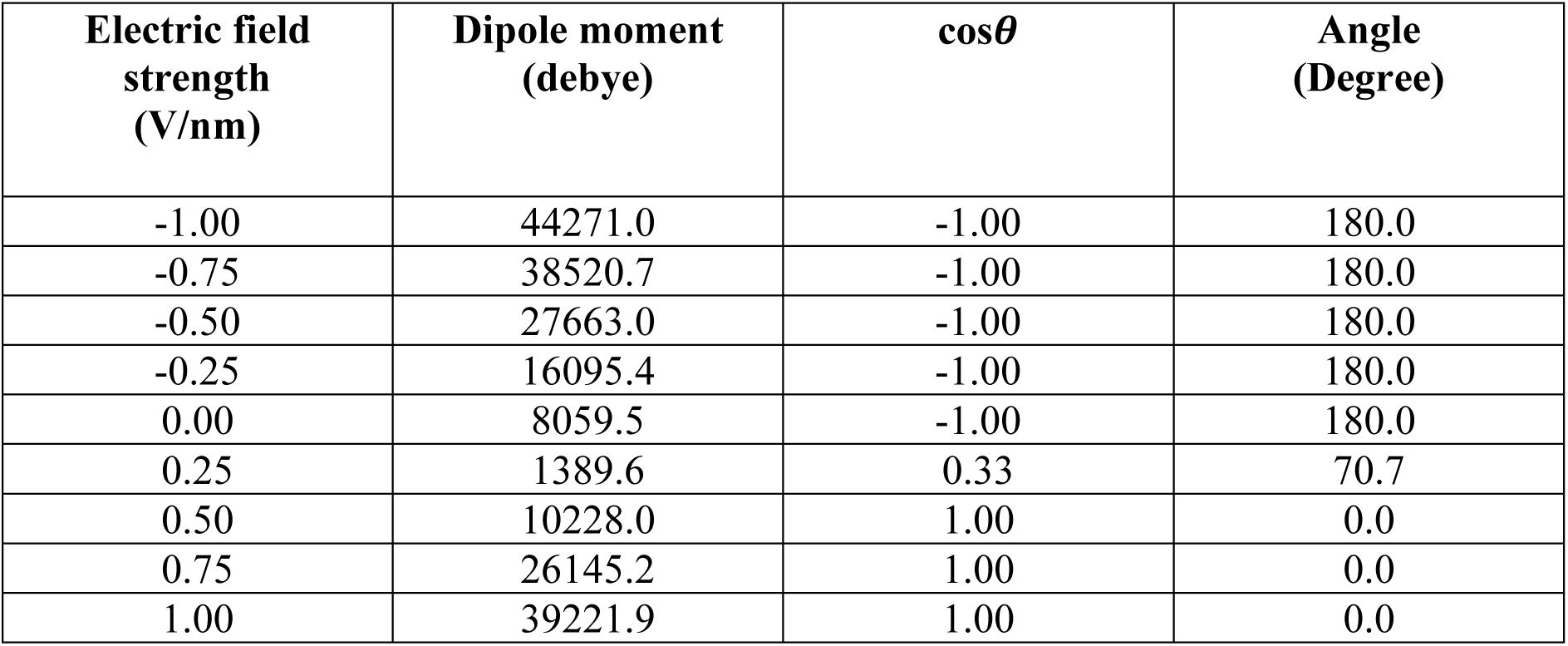
Dependency of magnitude and orientation of total dipole moment vector of water on applied electric field. Value presented here is the instantaneous magnitude of dipole moment vector at 100 ns.

The relative orientation of dipole moment vector of fibronectin, HA and water has been presented in Fig. 8 together with the principal axes of moments of FN. With the increase of field strength, all dipole moment vectors tend to align themselves parallel to the field direction (Fig. 8). The interaction among the dipole moment vectors of protein, HA and water acts as a major driving force of protein adsorption process, as described in the next section.

**Fig. 8:**
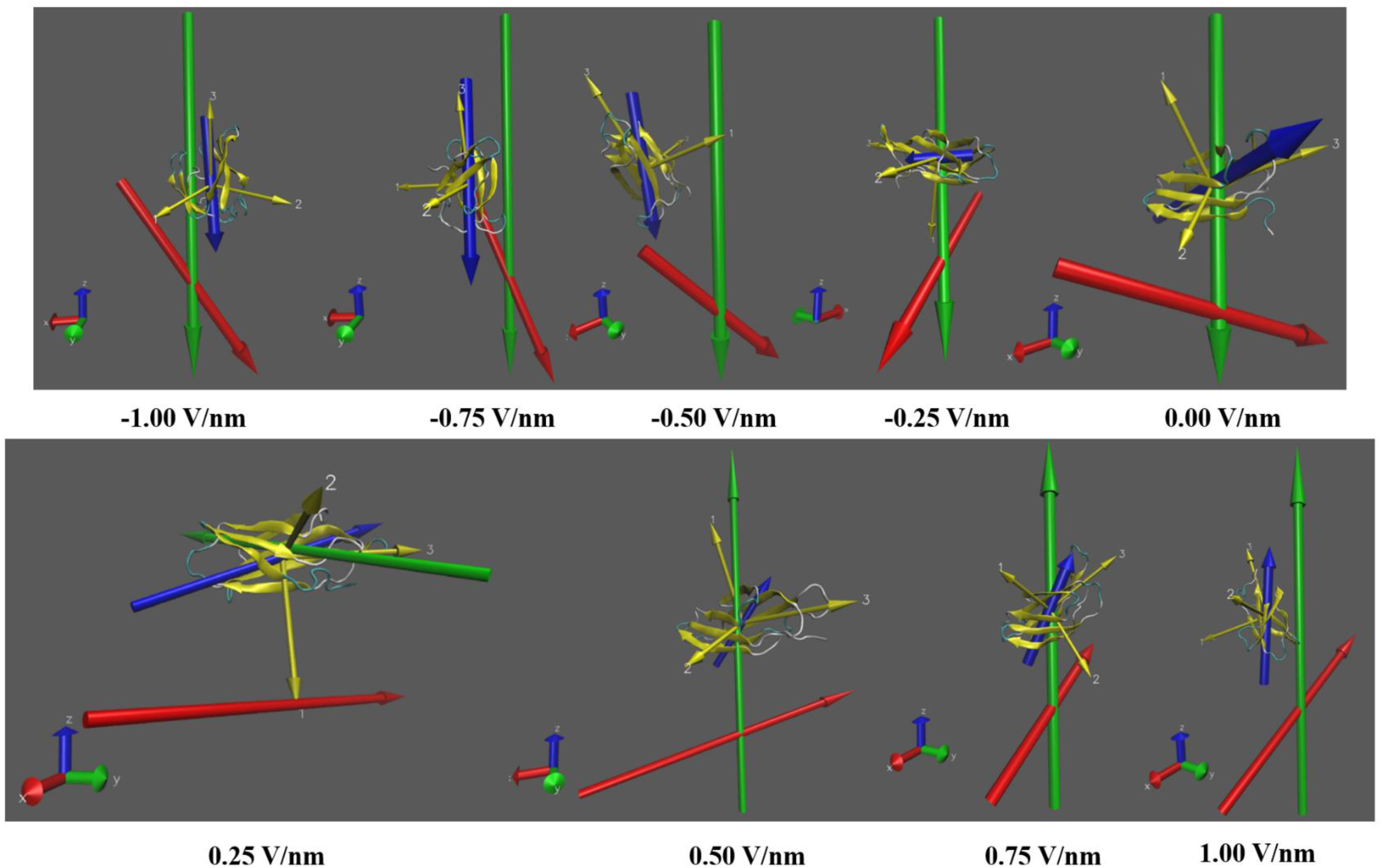
Dipole moment vector spontaneously align with electric field direction. Dipole moment orientation of protein (blue), HA (red) and water (green) at different field strengths and directions. The principal axes of momenta of FN is shown in yellow. At high field strength, the dipole moment vectors of FN and water became parallel to each other, which increased the mutual repulsion between them and affected the adsorption process (Axis colour code: Red: x, green: y, blue: z). Length of the arrows presenting dipole moment vectors are not to scale.

### 3.4 Solvation shell around fibronectin

The hydration shell around the protein and the surface plays an important role in adsorption phenomena. In order to get an insight into the water structure surrounding the protein, we have also calculated solvent radial distribution function using gmx tools. Fig. S11 in supplementary section presents *g*_*OC*_*(r)* of the water oxygen atoms surrounding C_α_ atom of a hydrophilic (Ser 53; Fig. S11(a)-(b)) and a hydrophobic (Leu8; Fig. S11(c)-(d)) residue at different electric field strengths. *r*_max_ was chosen as 2 nm. For both hydrophilic and hydrophobic residue, the radius of the first hydration shell was noted to be ∼0.4 nm, irrespective of the field strength /direction (Fig. S11 in supplementary section). For hydrophilic residue, the amplitude of the first peak of *g*_*OC*_*(r)* appeared almost identical except for high field strength, where it decreased. This observation implies less presence of solvent surrounding the reference atom. On the other side, almost no change has been recorded for *g*_*OC*_*(r)* of the hydrophobic residue, except at -1.00 V/nm (Fig.S11 in supplementary section). The significance of obtained results have been included in the supplementary information (section S3 of SI).

## 4. Discussion

In general, proteins change their conformation during the adsorption process, which depends on the substrate properties and the nature of the protein solution.^35, 36, 37^ Those conformational changes have grave influence on the cell-material interaction. For instance, bacteria exhibit different affinity towards saliva protein adsorbed on a material surface, compared to that in aqueous solution.^38^ Also denatured protein cannot support cell adhesion on material substrate. It is noteworthy that, not all conformational changes are beneficial for cell attachment. Hence, it is important to explore conformational changes of proteins for better understanding of biological response of materials.

Different experimental techniques like atomic force microscopy (AFM), time of flight secondary mass spectroscopy (ToF-SIMS), Fourier transform infrared spectroscopy (FTIR), etc. have been used to probe the conformational changes of proteins during adsorption.^39, 40, 41, 42^ However, probing protein material interaction at single molecular level is extremely difficult by experimental means and the nature of the interactions are not completely known till this date.^43^ Computational techniques come to rescue as they offer to explore the area beyond the experimental regime. MD simulation is arguably the most widely used approach to investigate interaction between proteins and inorganic surfaces. Different types of surfaces, namely pure metals, metal oxides, carbon allotropes, have been modelled to study their affinity towards proteins.^44^ Among them, hydroxyapatite (HA), being the hand tissue mineral is of particular interest because of its proven biocompatibility in living systems. Between two stable crystallographic structures of HA, hexagonal structure is widely recognized due to its presence in hard tissue.45 Hexagonal HA exhibits P6_3_/m crystallographic symmetry with the lattice parameters: *a=*9.432 Å and *c=*6.881Å.^46^

In this section, a correlation among different results penned in the previous section will be drawn together with the detailed analysis of the underlying causes behind the recorded behaviors of proteins. In order to achieve this, the established mechanisms of protein-hydroxyapatite interaction will be reiterated first.

### 4.1 Influence of electrostatic interaction energy on the adsorption kinetics

As mentioned before, the adsorption of FN on the HA surface is mediated by electrostatic interaction. It is noteworthy that not all protein residues interact equally with the HA surface. Rather, there exists specific binding sites for this particular interaction. Phosphorylated amino acids constitute the first class of HA binding sites and the second class is made up of γ-carboxyglutamic acid, found in the mineral binding portion of bone Gla protein.^47^ While these two classes are composed of modified amino acids, a third class of HA binding site comprised of unmodified amino acids.^47^ The candidate for the third class of HA binding site is mainly the sequence of acidic amino acids.^47^ For example, Asp or Glu-rich sections of proteins are found to bind with the HA crystal.^47^

Amino acids mainly interact with HA through the formation of hydrogen bonds and electrostatic forces.^43^ In case of zwitterionic amino acids, interaction takes place via Ca-O bond formation. Due to this fact, many amino acids tend to adsorb on HA surface in their zwitterionic form.^48^ An oft-called example is the adsorption of Gly on the HA (010) surface, as iterated in the previous section.^11^ The formation of Ca-O and O-Ca-O bonds through proton transfer from amino acid to HA was evident from DFT calculation.^11^ Similar adsorption phenomena were recorded in the case of Lys.^7^ Apart from the bond formation, electrostatic interaction between the oppositely charged groups of protein and HA (COO^-^ and Ca^2+^ ; NH_3_^+^ and PO _4_^3-^) also dominates the protein adsorption, as noticed in many cases.^48, 16^ Another important mode of interaction is water bridge formation between protein and HA. In the case of BMP-2 adsorption on HA (001) surface, water bridged H-bonds were found to dominate the attractive interaction.^49^

In the present study, the electrostatic interaction was found to mediate the adsorption phenomena. Mainly the positively and negatively charged amino acids interacted with the HA surface (Table 2). As mentioned before, the positively and negatively charged residues interact with the surface via their positively and negatively charged groups, respectively. Clearly, the adsorption was driven by the electrostatic interaction among the oppositely charged groups of FN and HA. In some cases, neutral residues came closer to the surface in the course of simulation, but their interaction with the surface was found to be weak compared to that of the charged residues (Fig. S12 in supplementary information).

### 4.2 Probing of E-field mediated protein conformal changes and stability

When a protein is exposed to the external electric field, two types of possible phenomena can take place in order to compensate the effect of the field. First, the rearrangement of the positions of the local atomic charges, and the global conformational change of the whole protein.^50^ The combinatorial effects of these two phenomena result in the recorded behavior of fibronectin on the HA surface. When electric field strength was 0.25 V/nm, the first phenomena dominated and the RMSD value got lower, compared to that in absence of the electric field. The local rearrangement of charges brought more residues near the HA surface (Table 2). As a consequence, the number of hydrogen bonds between HA and FN increased together with the attractive electrostatic energy (Fig. S2 in supplementary section). Hence, a strong adsorbed state was observed along with reduced FN-HA distance (Fig. 1).

When the field strength was increased to 0.50 V/nm, global structural changes of FN began to take place together with the local rearrangement. This was reflected in the increased RMSD value and reduced number of interacting residues (Fig. 6 and Table 2). Due to the presence of a smaller number of residues near the surface (Table 2), the number of hydrogen bonds and atomic contacts also got declined. The attractive interaction energy also decreased, and FN-HA distance increased. As a result, the adsorbed state weakened. The similar trend followed with further increment of the field strength. Hence, one can conclude that, beyond 0.25 V/nm, the application of electric field can not support FN adsorption on the HA surface.

The scenario was a bit different, when the field direction was reversed. At the field strength of -0.25 V/nm, the number of interacting residues decreased compared to 0.00 V/nm, leading to a decline in hydrogen bond formation and number of contacts (Fig. S2 in supplementary section and Fig. 3). There was also a little drop in average attractive interaction energy (Table 1). This is probably due to the fact, rearrangement of positions of local atomic charges and global structural changes of FN happened in a different manner on the HA surface. The similar trend followed at field strength of -0.50 V/nm. Here, the changes of the whole FN module were more prominent, as evident from the higher RMSD. The adsorption was also weak at this field strength. Only a few residues interact with the HA (001) surface (Table 2). As these residues are part of the RGD loop, the application of high electric field in an antiparallel direction is not suitable for integrin-FN interaction. Hence, cell-material interaction will not be facilitated in the presence of anti-parallel field direction.

When field strength was further increased to -0.75 V/nm, similar behavior like -0.50 V/nm, was observed. The interacting residue was same as -0.50 V/nm. The notable difference is that the attractive interaction energy, number of contacts and the frame average number of hydrogen bonds, both increased at -0.75 V/nm field strength (Fig. 2, 3 and S3 in supplementary information). The underlying reason may be the different structural changes of FN under the influence of these two electric field strengths. Moreover, the main contribution towards the interaction energy and hydrogen bond formation came from the interacting Arg78 residue. It has been observed that, between ∼54 ns and ∼90 ns, the COM-COM distance between Arg78 and HA was higher in case of -0.50 V/nm, compared to that at -0.75 V/nm (Fig. S14(a) in supplementary information). No hydrogen bond was formed between Arg78 and HA during that time period (Fig. S14(b), supplementary information). This has been reflected in the average number of H-bonds. Due to similar reason, Arg78 interacted with HA more strongly in case of -0.75 V/nm (Fig. S14(c) in supplementary section) and total interaction energy increased (Fig. 2).

At field strength -1.00 V/nm, the protein-FN distance and RMSD increased significantly. Only two residues were close to the surface (Table 2) and the neutral residue interacted weakly with the HA surface (Fig. S12(c), supplementary). The major contribution towards interaction energy and hydrogen bond formation came from the interaction of Val1 residue with HA (Fig. S13, supplementary information). The number of absorbing residues increased at -1.00 V/nm, and this was reflected in the slight increment of number of contacts (Table 2 and Fig. 3).

The prolate shape of FN was confirmed in absence of HA surface from the ratio of I_1_/I_3_ (Table 3). The same shape was retained in presence of surface and electric field, as well. As the ratio of I_1_/I_2_ was found to be close to 1, the ellipsoid shape of FN can crudely be approximated as a prolate spheroid, in all cases. From the values of principal components of moment, it can be concluded that, FN first got elongated along the *c* axis of the ellipsoid up to +0.50 V/nm and thereafter, it got shrank. The same pattern was followed for negative field direction, as well. However, in this case, the decrease of *c* axis started after -0.25 V/nm (Table 3).

There exists a clear dependency of protein behavior on the field direction. The adsorption of FN was noted to be weaker when the field was applied antiparallel to the z-axis. Moreover, the RMSD was also found to be higher. The probable reasons can be the dipole-dipole interaction among FN, HA and water. As mentioned before, the dipole moments of water, protein and HA tended to align themselves parallel to the field direction. It is a well-known fact that the parallel dipoles experience repulsive force and anti-parallel dipoles experience attractive force. Among several factors, the force between dipoles depends on the magnitude of dipoles and on the cosine of the angle between them.^51^ In particular, with the increase of the angle (upto 180°), the repulsive nature decreases. In the present case, as the field strength was increased, the angle between dipole moment vector of water and FN decreased, irrespective of field direction. As a consequence, the repulsive force between the dipoles increased, resulting in a weak adsorbed state at high electric field strength. Moreover, when field was negative, the magnitude of dipole moments for both FN and water was higher compared to their counterparts with positive electric field. This led to a comparatively weaker adsorbed states of FN at negative electric fields.

The other important aspect of our study is to analyze the stability of the FN in the presence of electric field. As mentioned before, the intramolecular salt bridge network helps in protein structure stabilization.^52^ In the current study, it has been found that, although more salt bridge networks were found to be formed at high electric field strength (Table 4), the overall structural integrity was affected in high field strength. The percentage of occurrence of different salt bridges also strongly depends on the electric field strength and the nature of dependency is specific to the residues taking part in the salt bridge formation (Fig. S7 in supplementary information). Besides this, the temporal stability of the salt bridges also reduced with the increase of the field strength, in general (Fig. S7 in supplementary information). All of these phenomena can be attributed to the local rearrangement of the atomic positions of protein together with the global structural changes to counteract the effect of the external field. The effect of the electric field on O-N atomic pair distance in different salt bridges significantly varies, because of the different locations of salt bridge forming residues within the protein structure (Fig. S6 in supplementary information). This results in the observed non distinctive nature of the dependency of occurrence frequency (percentage of occurrence) on field strength and direction. At very high field strength, the structure of FN significantly varies from its initial structure (as evident from higher RMSD), affecting the stability of most of the salt bridges and thus, affecting the overall structural stability of FN.

## 5. Conclusions

In the present study, we have examined the effect of external field on the fibronectin adsorption on the HA (001) surface. The following conclusions can be drawn from the present study.

a. Adsorption of FN on HA can be modulated by means of external electric field. Adsorption was favoured at low field strength (up to +0.25 V/nm). Weakened adsorbed state was observed at high field intensity.
b. Fibronectin interacts with HA via electrostatic force. The charged residues contribute majorly in the FN-HA interaction. The average attraction interaction energy first increased (up to 0.25 V/nm) and then decreased with the increment of electric field (from -454.3 kJ/mol at 0.00 V/nm to -245.2 kJ/mol at 1.00 V/nm).
c. The biophysical separation distance between FN and HA surface increased from ∼32 Å (0 V/nm) to ∼42 Å (1 V/nm) and ∼66 Å (−1 V/nm). Such large separation distance indicates weaker adsorbed state at high field strength. Although at low field strength (upto +0.25 V/nm), adsorption was promoted.
d. Non-monotonous behavior of RMSD can be explained in terms of two underlying phenomena, which include spatial rearrangement of local atomic charges and global structural change of protein.
e. An application of high electric field in antiparallel to z-axis does not facilitate cell-material interaction due to the proximity of the RGD loop to the HA surface.
f. The weakening effect of external electric field on adsorption phenomena is more profound at negative field. This can be implicated to the dipole-dipole interaction between FN and water.

In summary, the current study elucidates the effects of electric field on the structural changes and adsorption of fibronectin on HA. This is of particular interest in the context of bioelectronics medicine, where FN adsorption plays an important part in the interaction between cellular and mineral part of hard tissue in the presence of electrical stimuli. The findings of our study will provide a primary guideline to optimize external stimuli-based design of bioelectronics devices.

## Acknowledgements

The authors acknowledge Sahasrat of Supercomputer Education and Research Center (SERC) and Thematic Unit of Excellence on Computational Materials Science (TUE-CMS) of Solid State and Structural Chemistry Unit (SSCU) at IISc, Bangalore for providing access to the high-performance supercomputer facilities. One of the authors (BB) acknowledge DBT-National Bioscience award for financial support. SB and BB also thank to Prof. P. Balaram and Prof. M.K. Mathew, NCBS, Bangalore for useful discussion. BG thanks to Dr. D. S. Kothari Postdoctoral Fellowship scheme by University Grants Commission (UGC), India, for financial support.

