## supporting figures and text for "Electric field mediated fibronectin-hydroxyapatite interaction: A molecular insight"

### Probing Molecular-level interactions into electric field mediated fibronectin-hydroxyapatite interaction

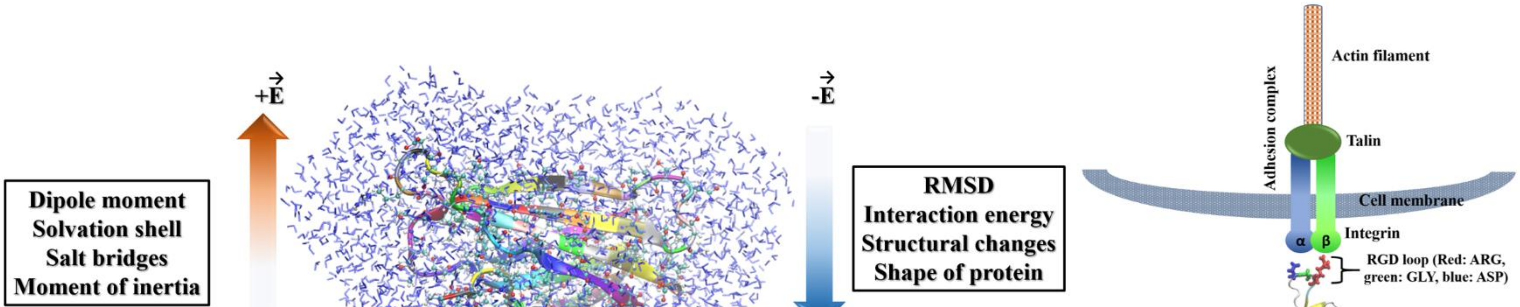

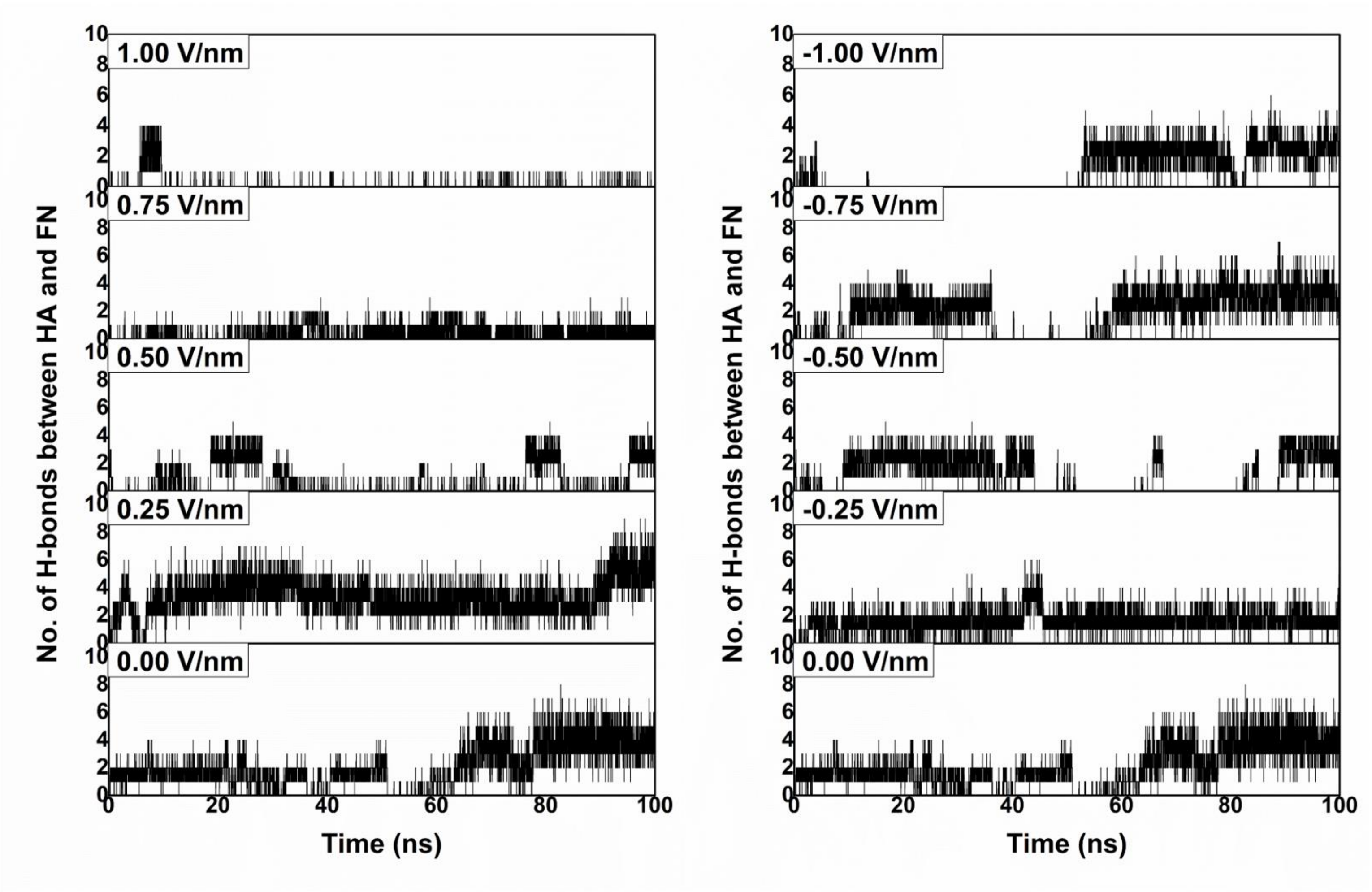


**Fig. S2: Formation of hydrogen bonds is one of the principal means of protein-material interaction.** Time-evolution of hydrogen bonds formed between fibronectin and HA at different field strengths.

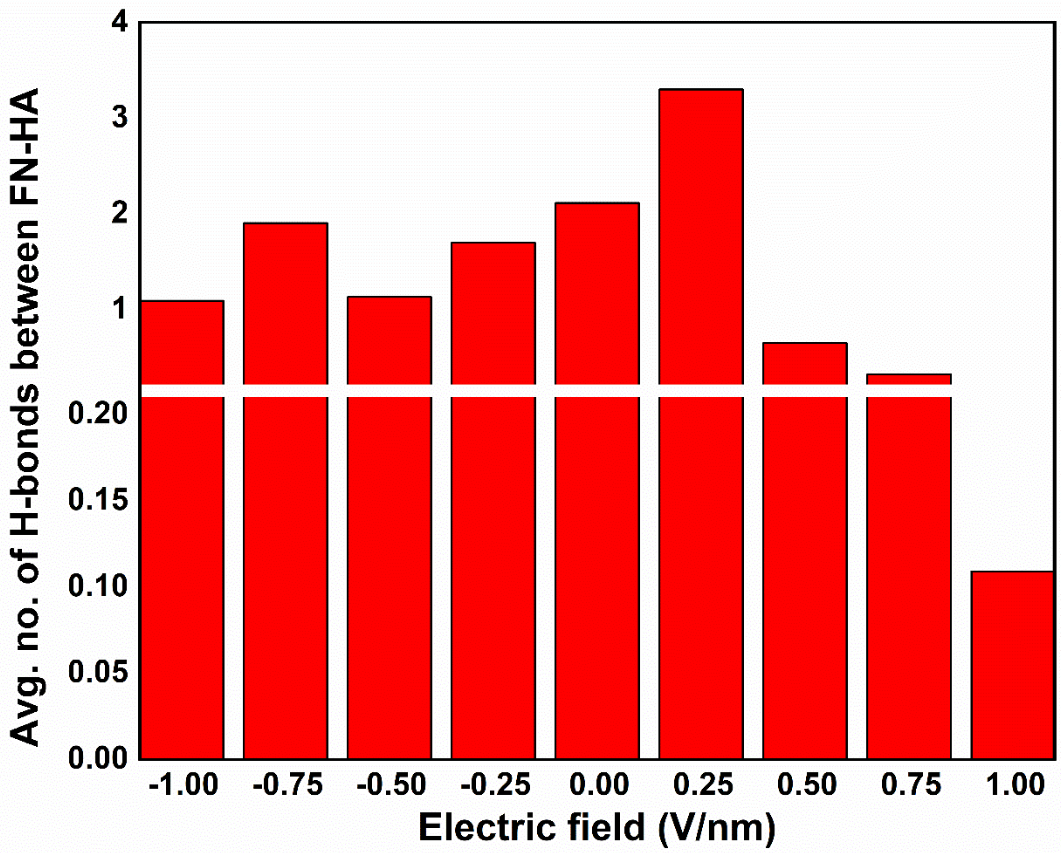


**Fig. S3:** Average number of hydrogen bonds formed between HA and FN at different electric fields.

#### S1. Hydrogen bond formation between FN and HA

Fig. S2 represents external field dependent time evolution of number of hydrogen bonds formed between HA and FN. The maximum number of hydrogen bonds was observed at field strength of 0.25 V/nm and these were reduced at higher field strength. The number of H-bonds was recorded to be comparatively higher at negative field values (Fig. S2). The decrease in the average number of hydrogen bonds was noted with an increase in the field strength, applied in an anti-parallel manner (Fig.S3). A slight increase was recorded at field strength of -0.75 V/nm (Fig. S3).

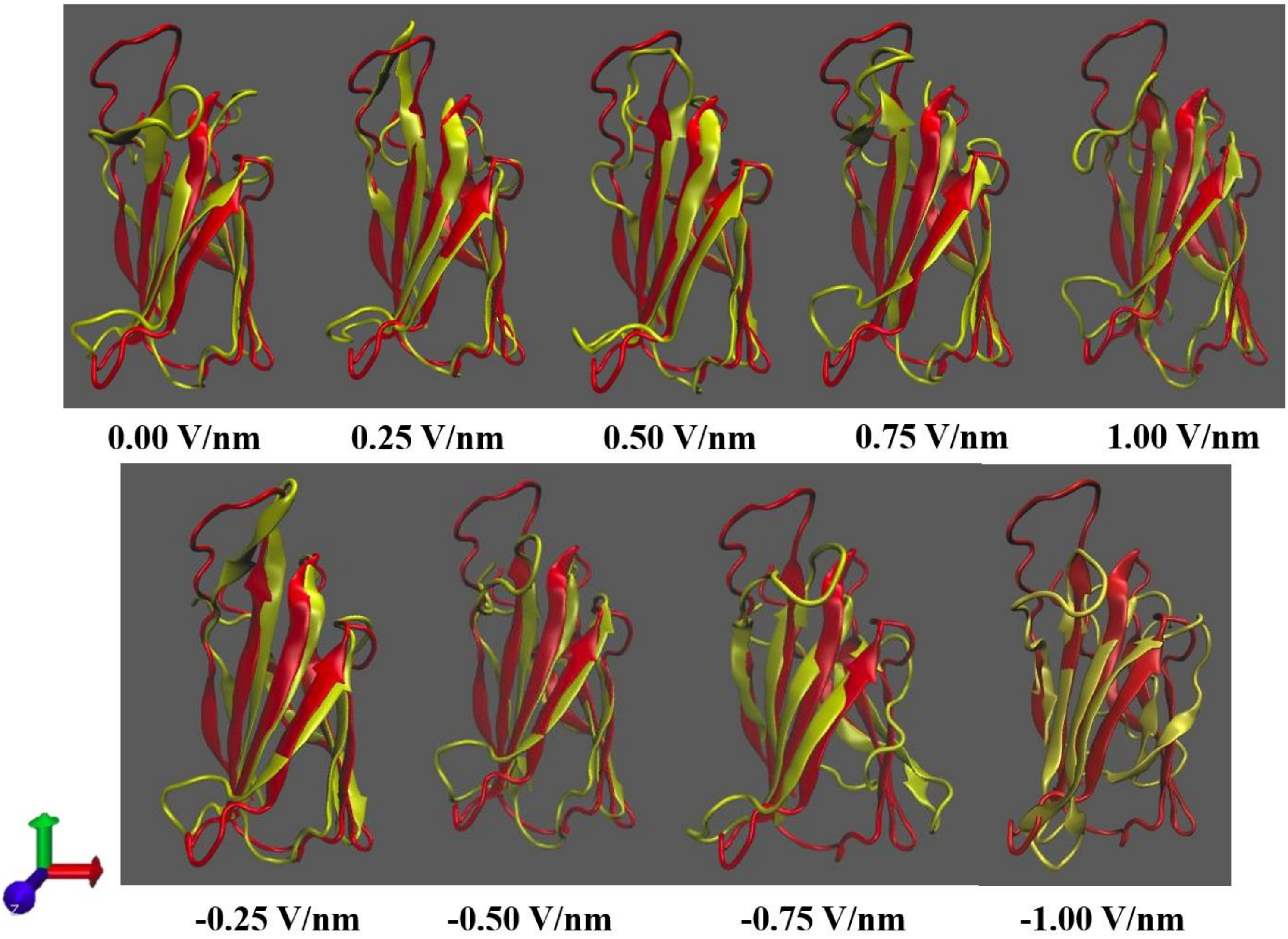


**Fig. S4: Electric field application can influence protein conformation.** Comparison of initial and final structure of fibronectin (colour code: Red: initial structure; yellow: final structure after 100 ns simulation. Axis colour code: Red: x, green: y, blue: z). Note that, conformational changes take place majorly in loop areas.

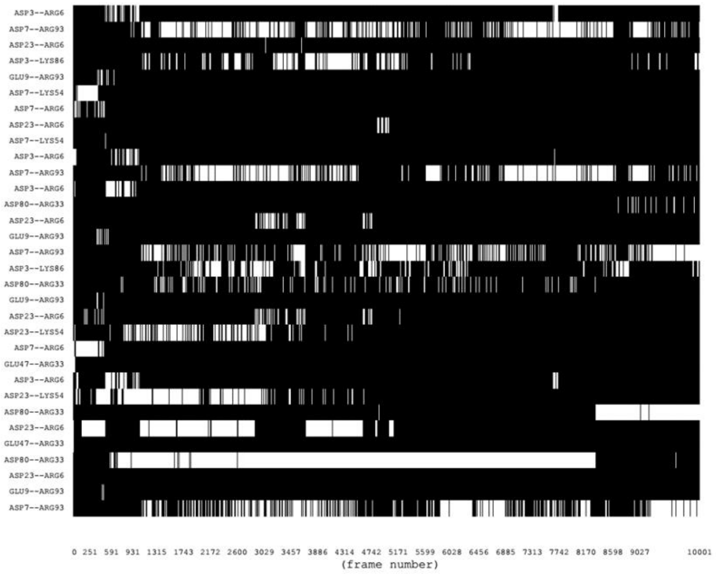


**-1.00 V/nm**

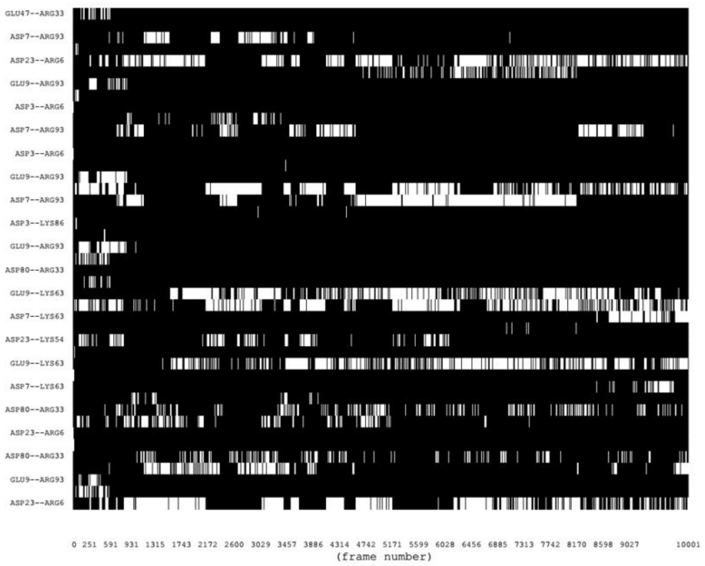


**-0.75 V/nm**

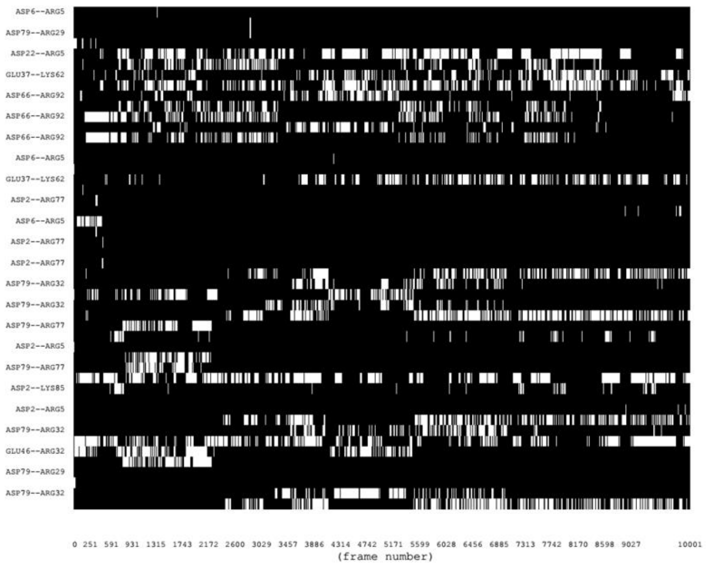


**-0.50 V/nm**

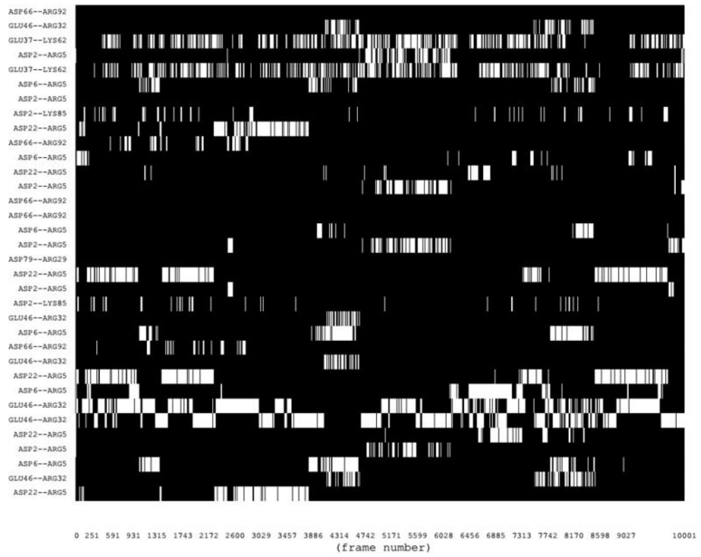


**-0.25 V/nm**

**(a)**

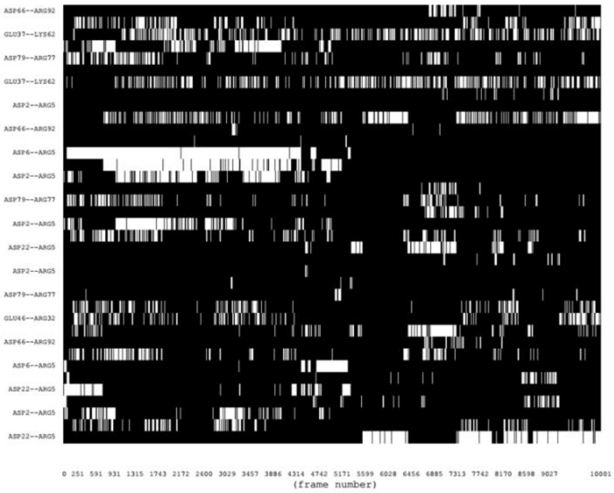

**(b)** **0.00 V/nm**

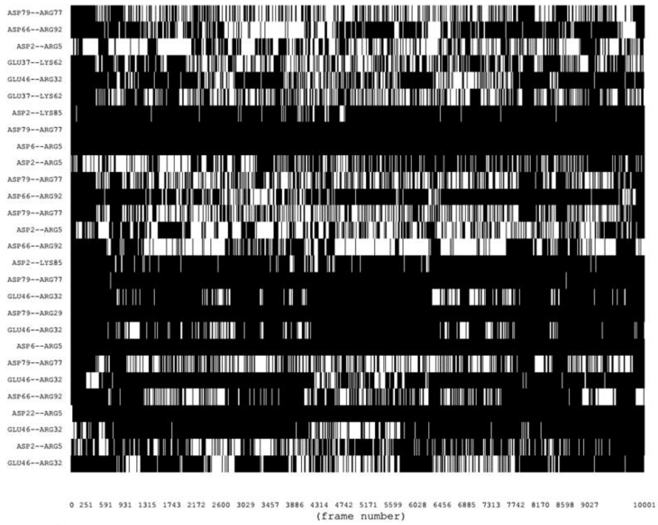

**0.25 V/nm**

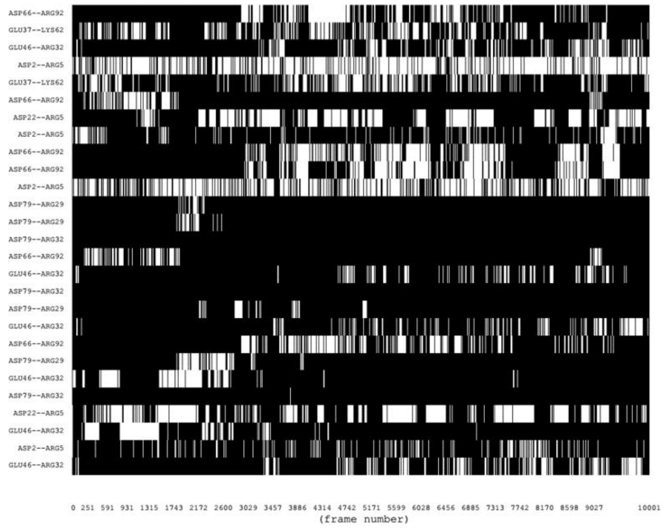

**0.50 V/nm**

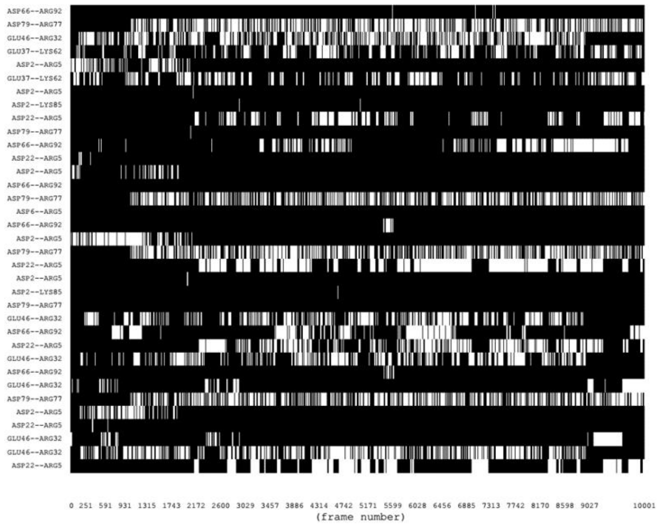

**0.75 V/nm**

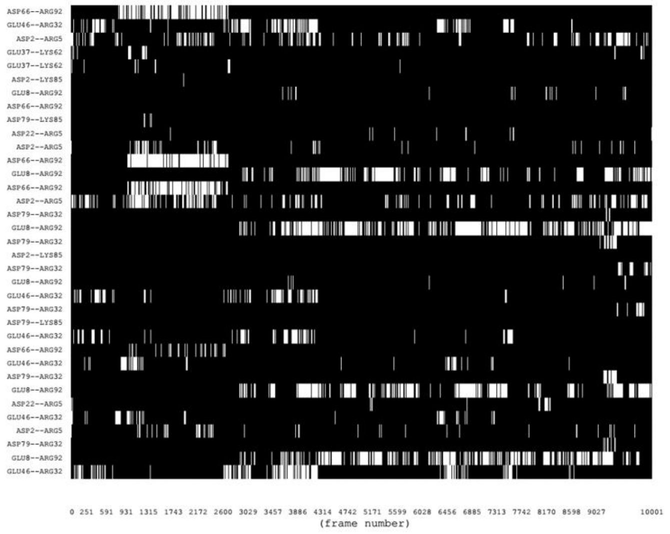

**1.00 V/nm**

**(c)**

**Fig. S5:** Temporal evolution of salt bridge networks of FN at (a) negative (b) zero and (c) positive electric fields. (colour Code: Black: no salt bridge, white: salt bridge present)

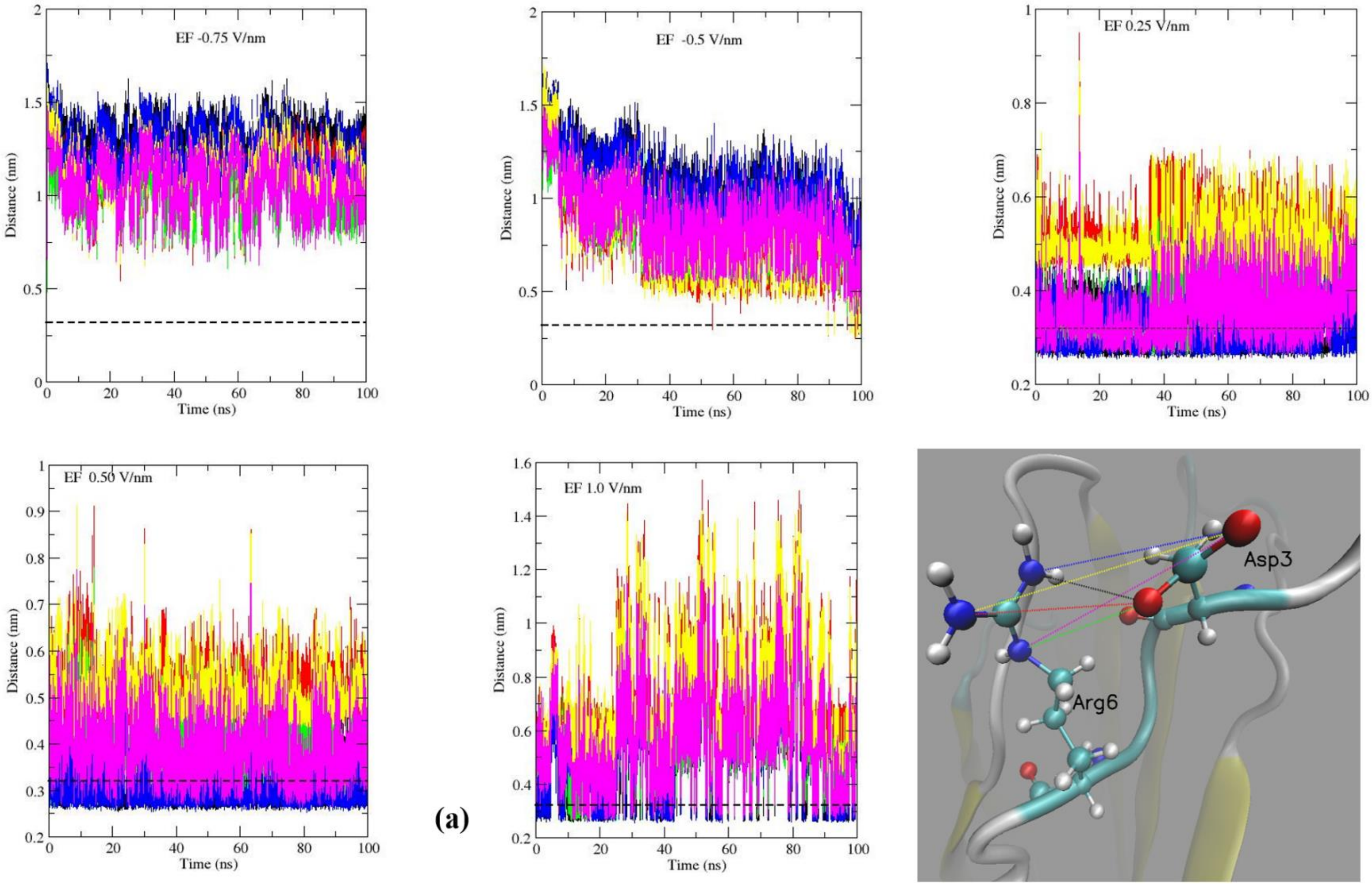
EF -0.75 V/nm
EF -0.5 V/nm
EF 0.25 V/nm
EF 0.50 V/nm
EF 1.0 V/nm
Distance (nm)
Time (ns)
Asp3
Arg6

(a)

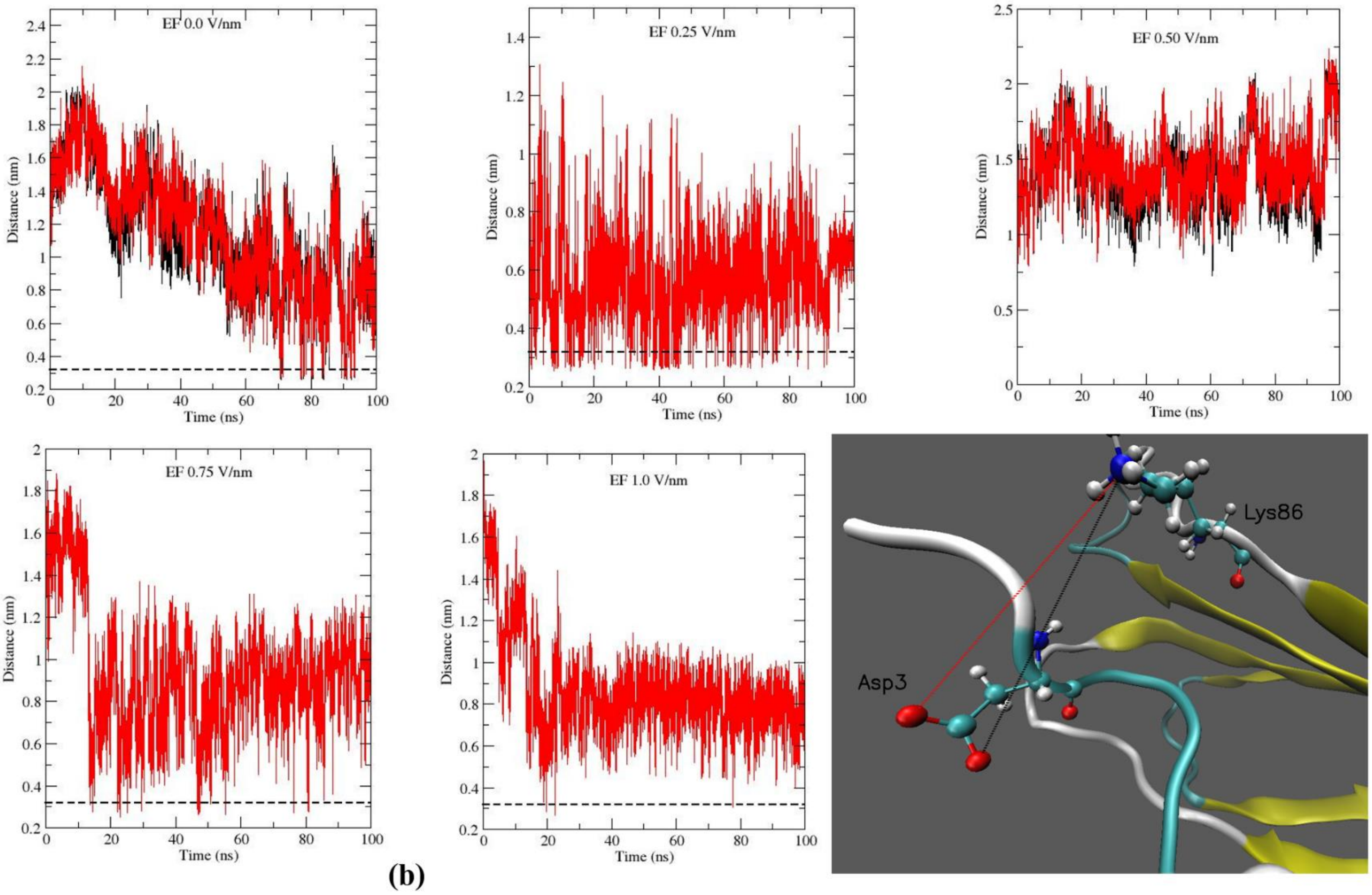
EF 0.0 V/nm
EF 0.25 V/nm
EF 0.50 V/nm
EF 0.75 V/nm
EF 1.0 V/nm
Distance (nm)
Time (ns)
Asp3
Lys86

**(b)**

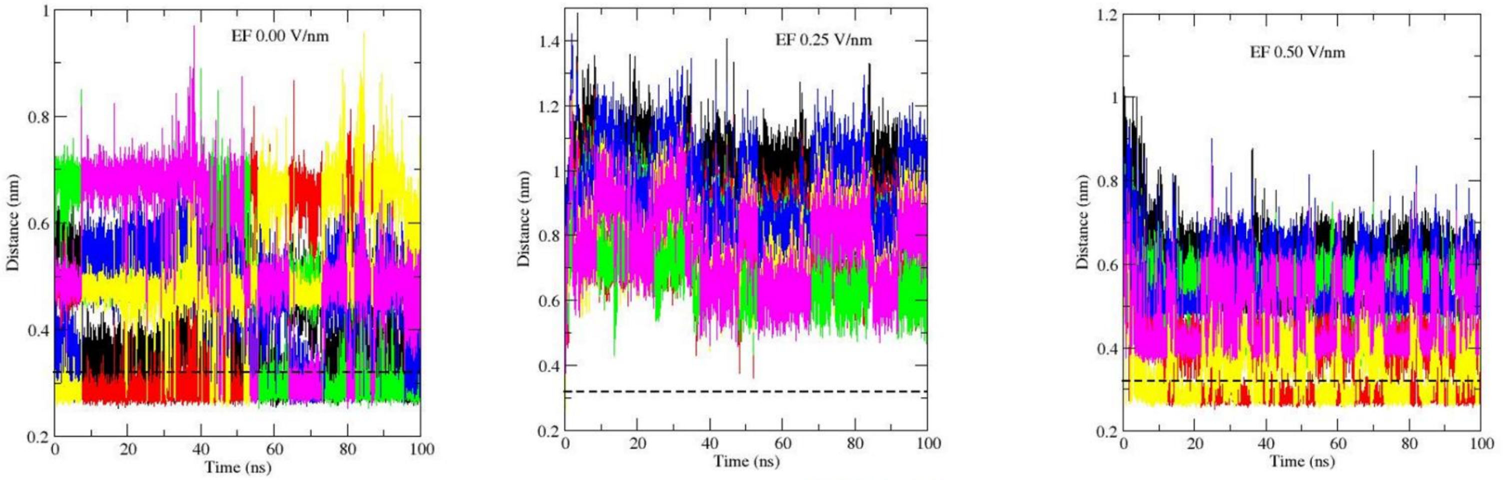
EF 0.00 V/nm
EF 0.25 V/nm
EF 0.50 V/nm
Distance (nm)
Time (ns)

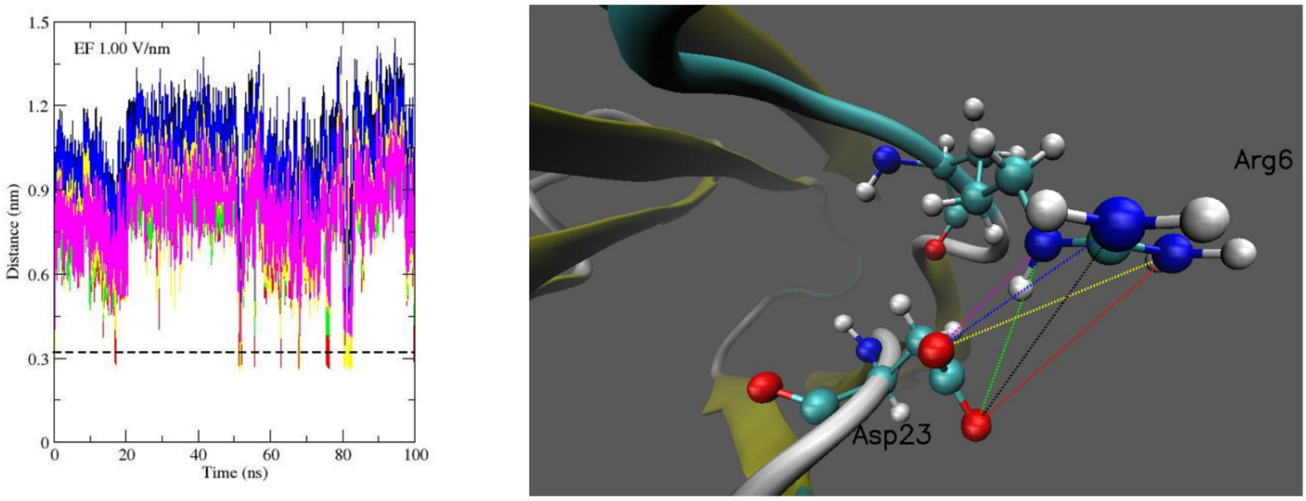
EF 1.00 V/nm
Distance (nm)
Time (ns)
Arg6
Asp23

**(c)**

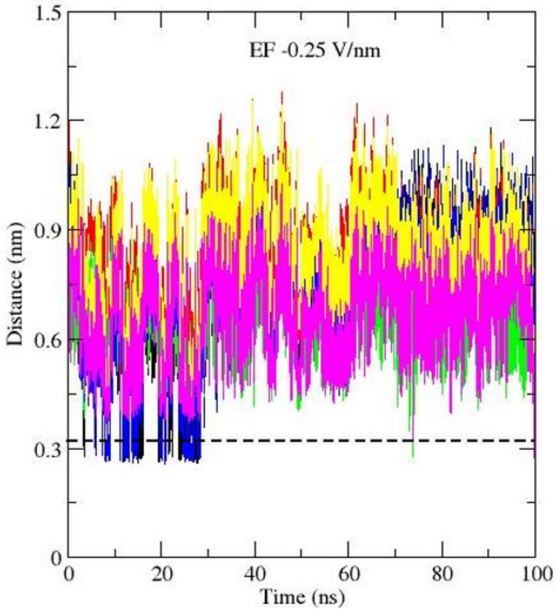
EF -0.25 V/nm
Distance (nm)
Time (ns)
1.5
1.2
0.9
0.6
0.3
0
0
20
40
60
80
100

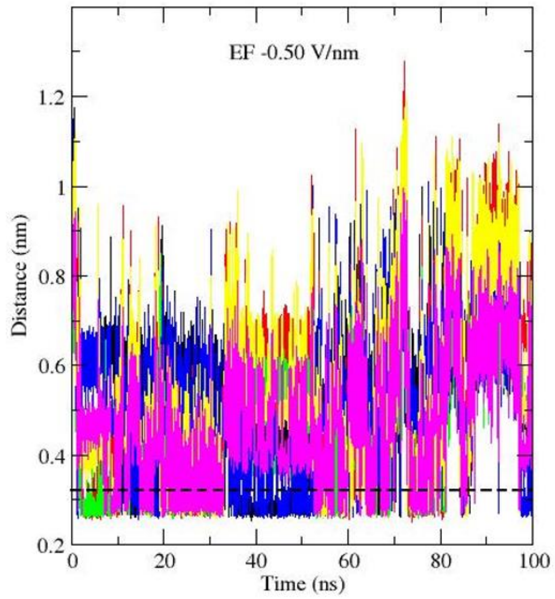
EF -0.50 V/nm
Distance (nm)
Time (ns)
1.2
1
0.8
0.6
0.4
0.2
0
20
40
60
80
100

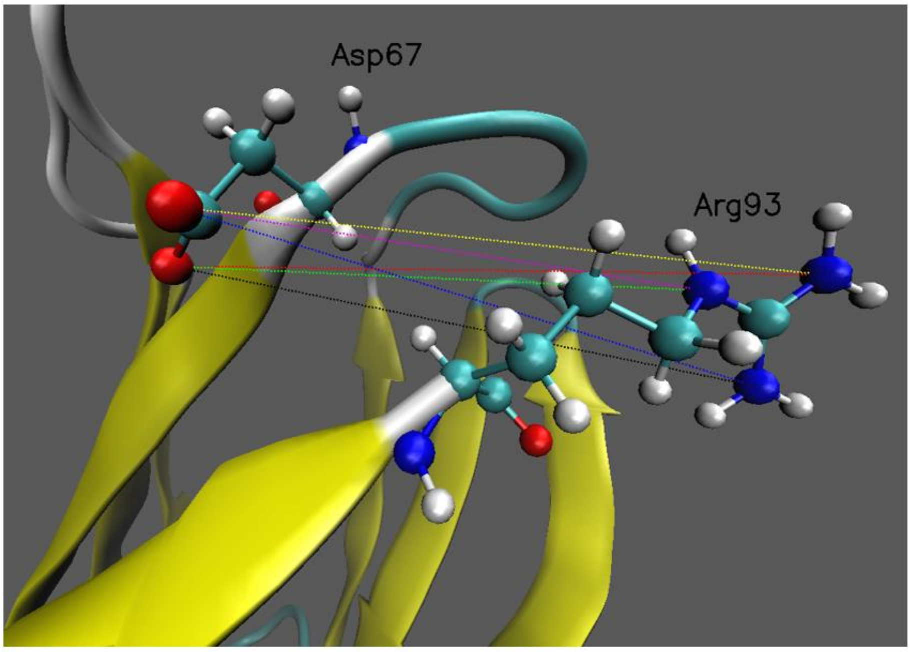
Asp67
Arg93

**(d)**

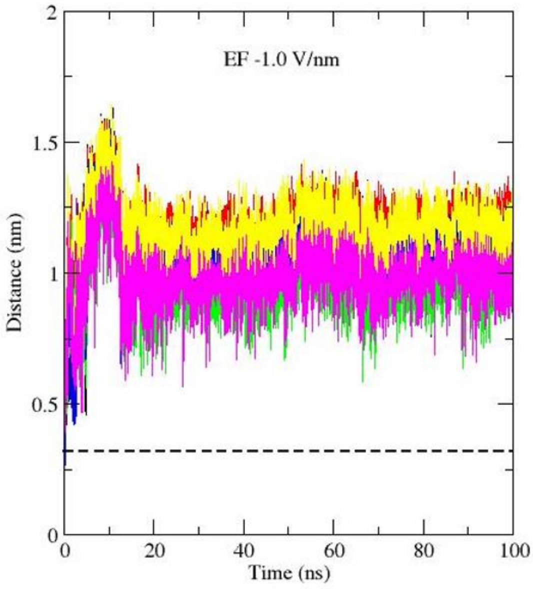


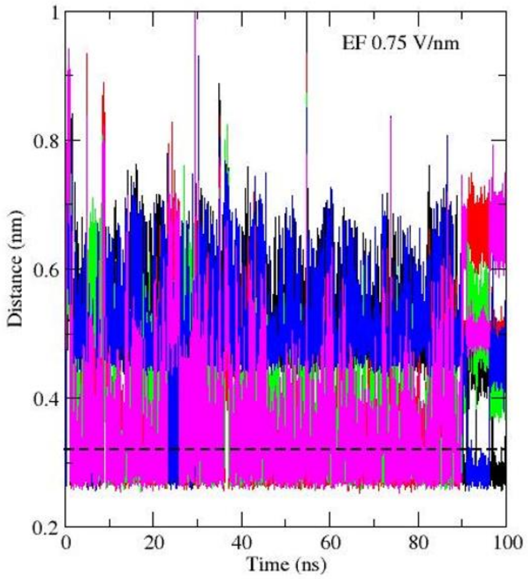


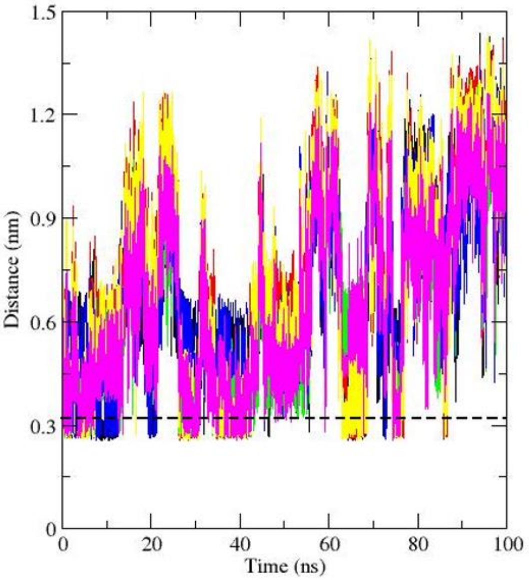


**(e)**

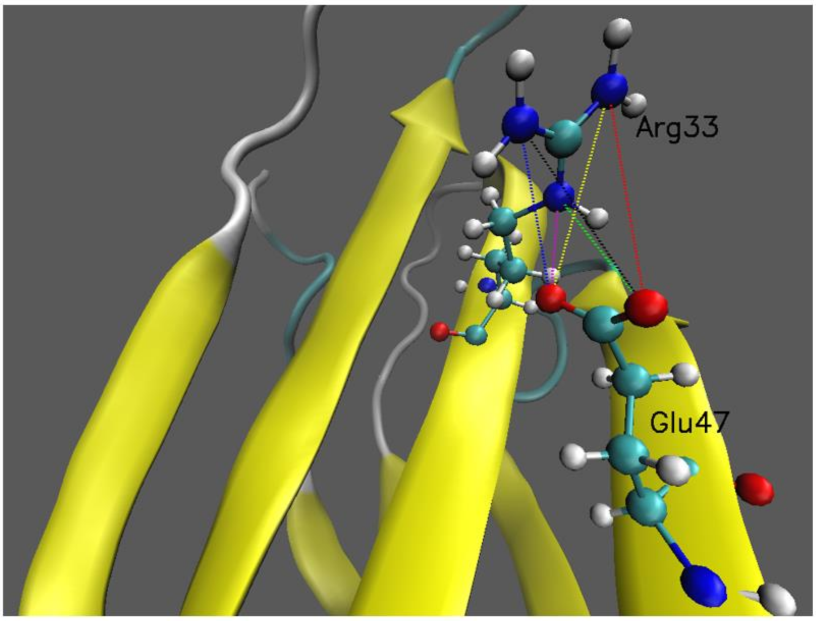


**Fig. S6:** Different salt bridges are switching on and off in the course of simulation. Distance between different O and N atomic pairs of (a) Asp3-Arg6 (b) Asp3-Lys86 (c) Asp23-Arg6 (d) Asp67-Arg93 and (e) Glu47-Arg33 salt bridge forming residues. Identical colours have been used to present the distances between different O-N atomic pairs in the plots and in the associated cartoons. Atomic colour code: Red: O, blue: N, cyan: C, white: H. The dashed horizontal line represents the cut off distance (0.32 nm) of salt bridge formation.

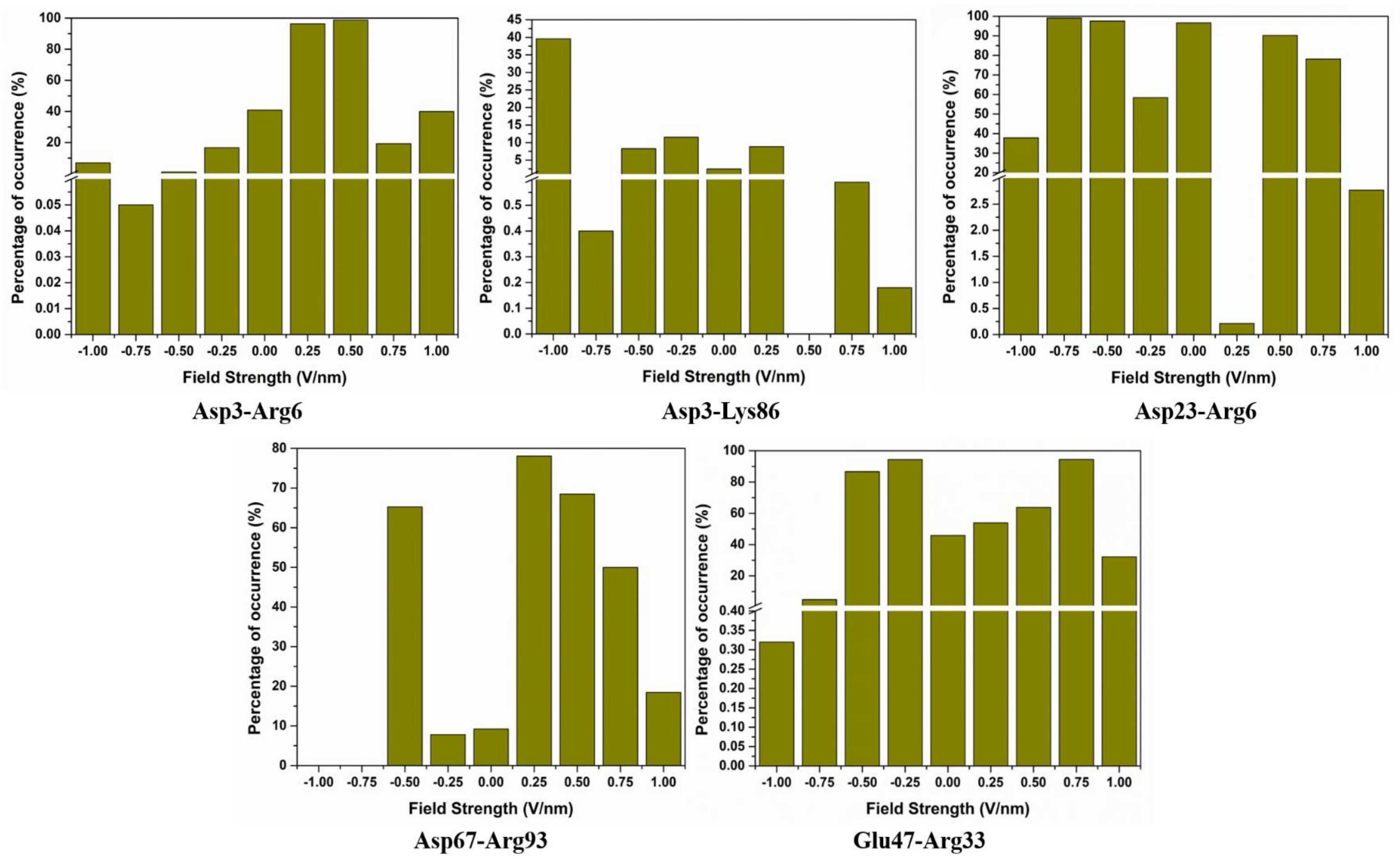


**Fig. S7: Salt bridges play important role in stabilizing protein structure.** Percentage of occurrence of different salt bridges at various electric field strength.

**S2. Temporal evolution of salt bridges**

The temporal evolution of different salt bridges has been presented in Fig. S5 in the supplementary section. A general observation is that the stability of the salt bridges was affected at high field strength (Fig. S5) In order to obtain a quantitative aspect, the percentage of occurrence (occurrence frequency) of some selected salt bridge nets were calculated and presented graphically in Fig.S7. From Fig. S7, it can be easily seen that different salt bridges responded differently under the influence of external field. The underlying reason is the non-identical behaviour of O-N atomic distances in different salt bridge forming residue pairs under the influence of same electric field intensity (Fig. S6). However, the occurrence frequency was found to be low at high field intensity in most of the cases, in corroboration with the information provided in Fig. S5 in supplementary section (Fig. S7). From this observation, it can be easily concluded that, the overall stability of the FN structure hindered at high field strength.

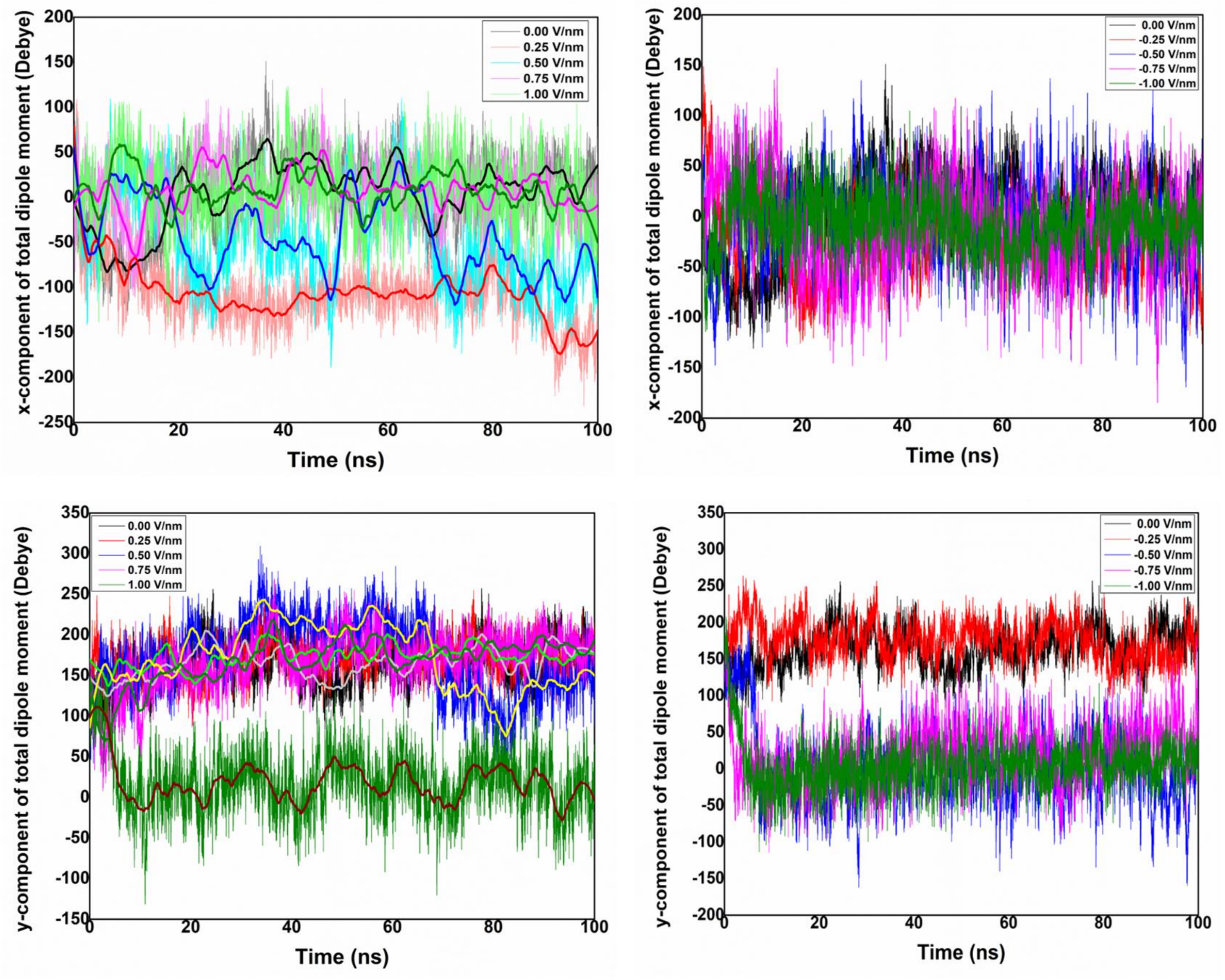


**Fig. S8:** x and y component of dipole moment of FN at different electric fields.

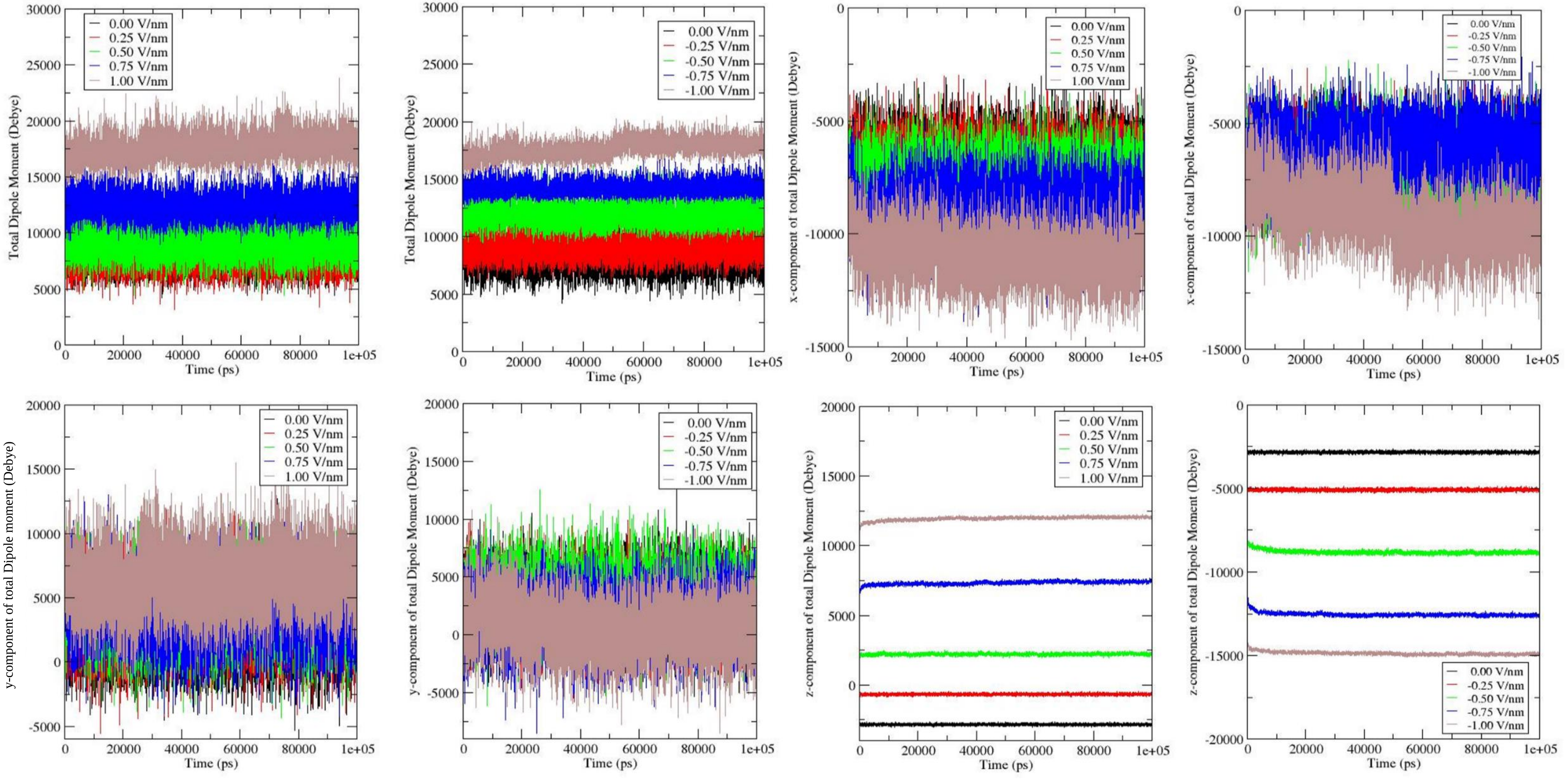


**Fig. S9:** Total, x, y and z component of dipole moment of HA at different electric fields.

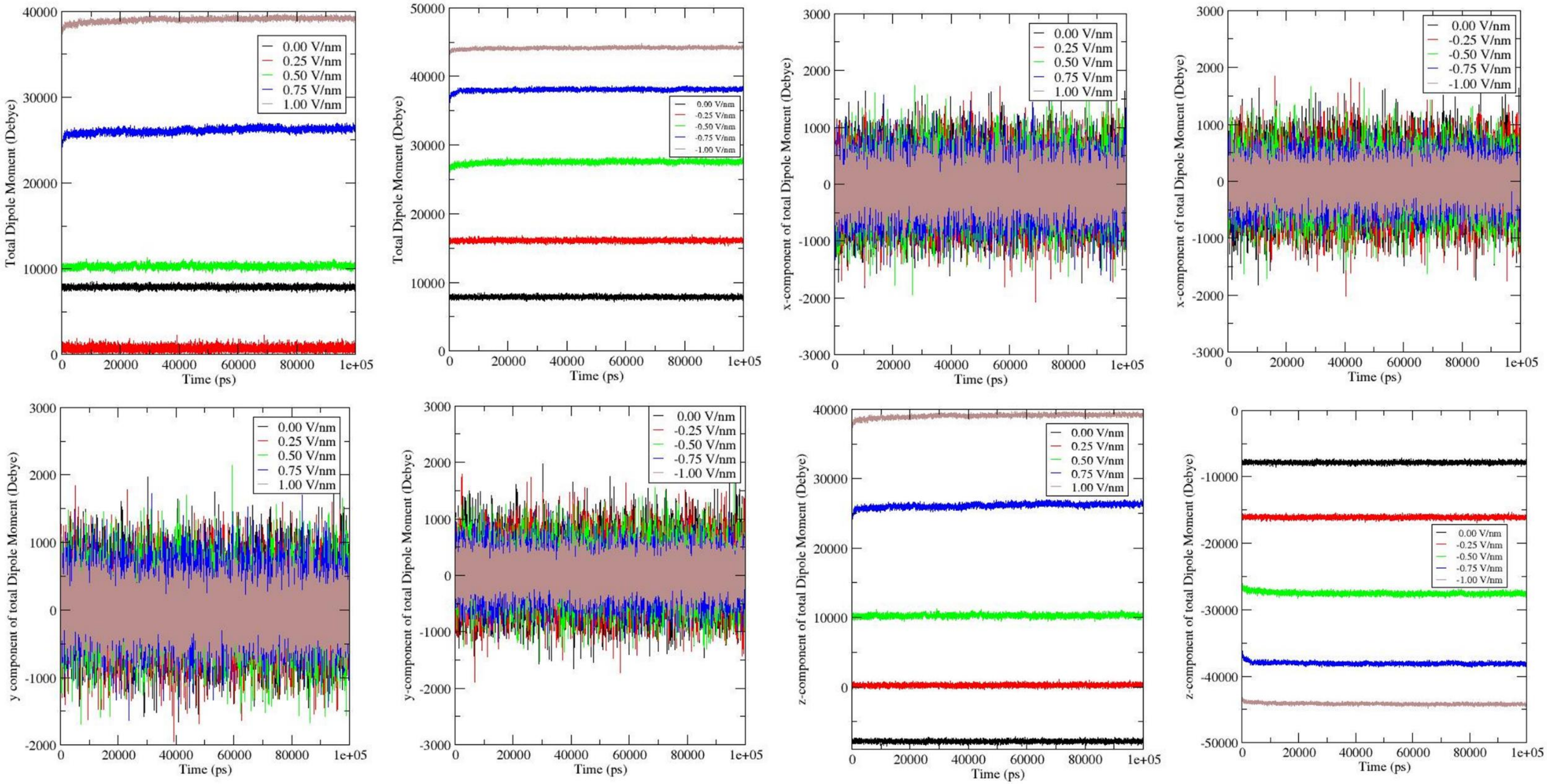


**Fig. S10:** Electric field dependency of total, x, y and z component of dipole moment of water.

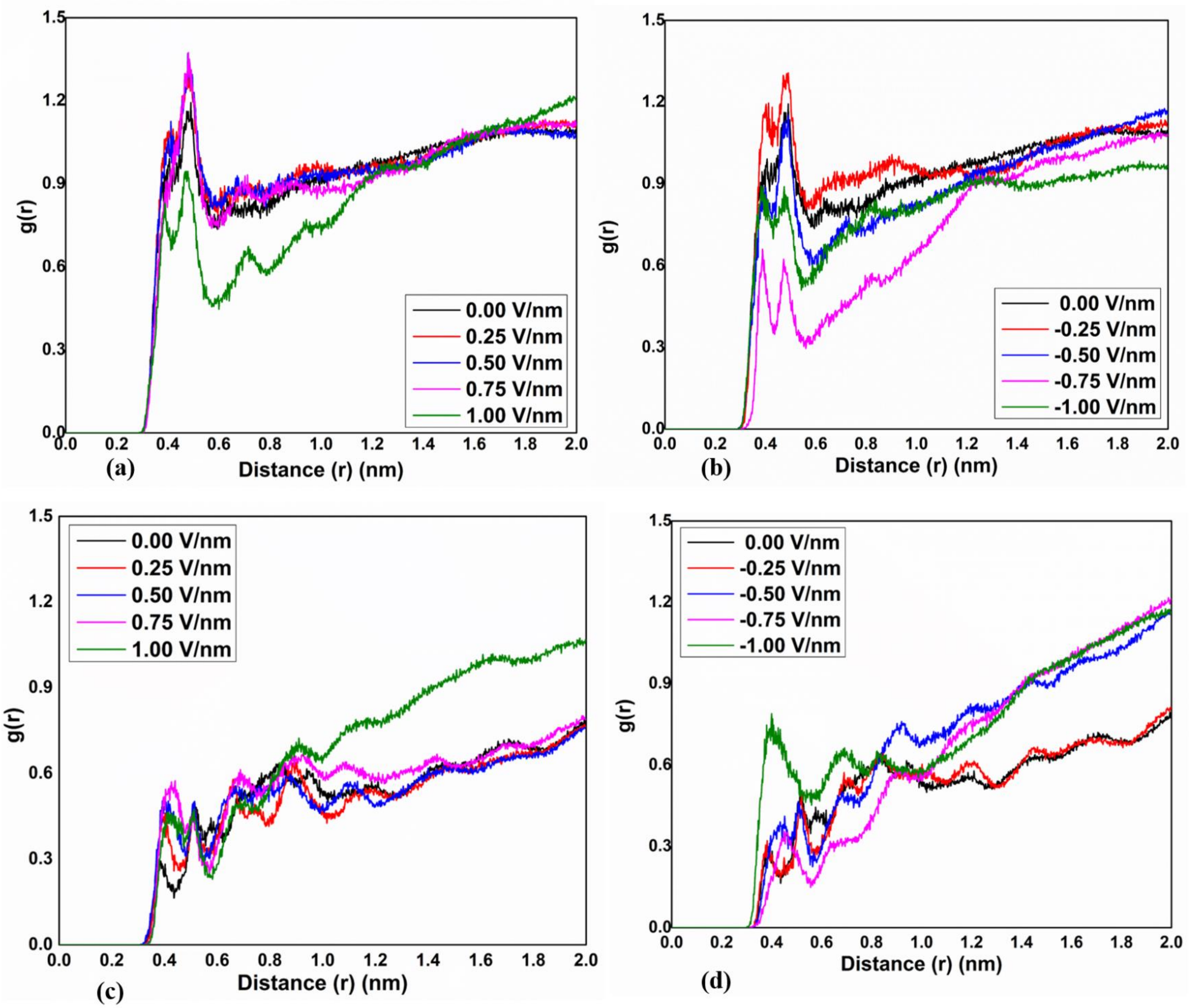


**Fig. S11: Solvation shell around the protein plays an important role in adsorption.** Radial distribution function (RDF; $g_{OC}(r)$) of water oxygen atoms surrounding $C_\alpha$ atom of a (a)-b) hydrophilic (Ser53) and (c)-(d) hydrophobic (Leu8) residue.

#### S3. Solvation shell around FN

As mentioned in the main article, the noticeable changes in the RDF peak intensity appeared only at high field strengths (Fig. S11). This indicates that, the presence of the electric field changed the solvation shell structure around the FN module, though not significantly.

There is a clear distinction between the arrangement of water molecules around the hydrophobic and hydrophilic residue (Fig. S11). The hydrophobic residues are usually buried into the inner surface of the protein and remains in limited contact with the solvent molecules.[1] On the other hand, the hydrophilic residues are exposed and are in direct contact with the water molecules.[1] This positional difference of the two types of residues results in the different orientation of water molecules surrounding them, which eventually leads to significantly different RDF. The intensity of the peaks decreased at high positive field strengths positive for hydrophilic residue probably, because of the structural rearrangement of FN (Fig. S11(a)-(b)). The structural rearrangement results in less accessibility of Ser53 for solvent molecules. Hence, the peak intensity decreased. A significant change in RDF of hydrophobic residue can be noticed at -1.00 V/nm (Fig. S11(d)). This observation can be corroborated with the high RMSD of FN at -1.00 V/nm (Fig. 6). At high field strength, the protein structure was found to be in its most altered condition and even the β-strands were affected (Fig. 6). Due to this conformal change, the inner part of FN got exposed to the solvent and the height RDF peak increased.

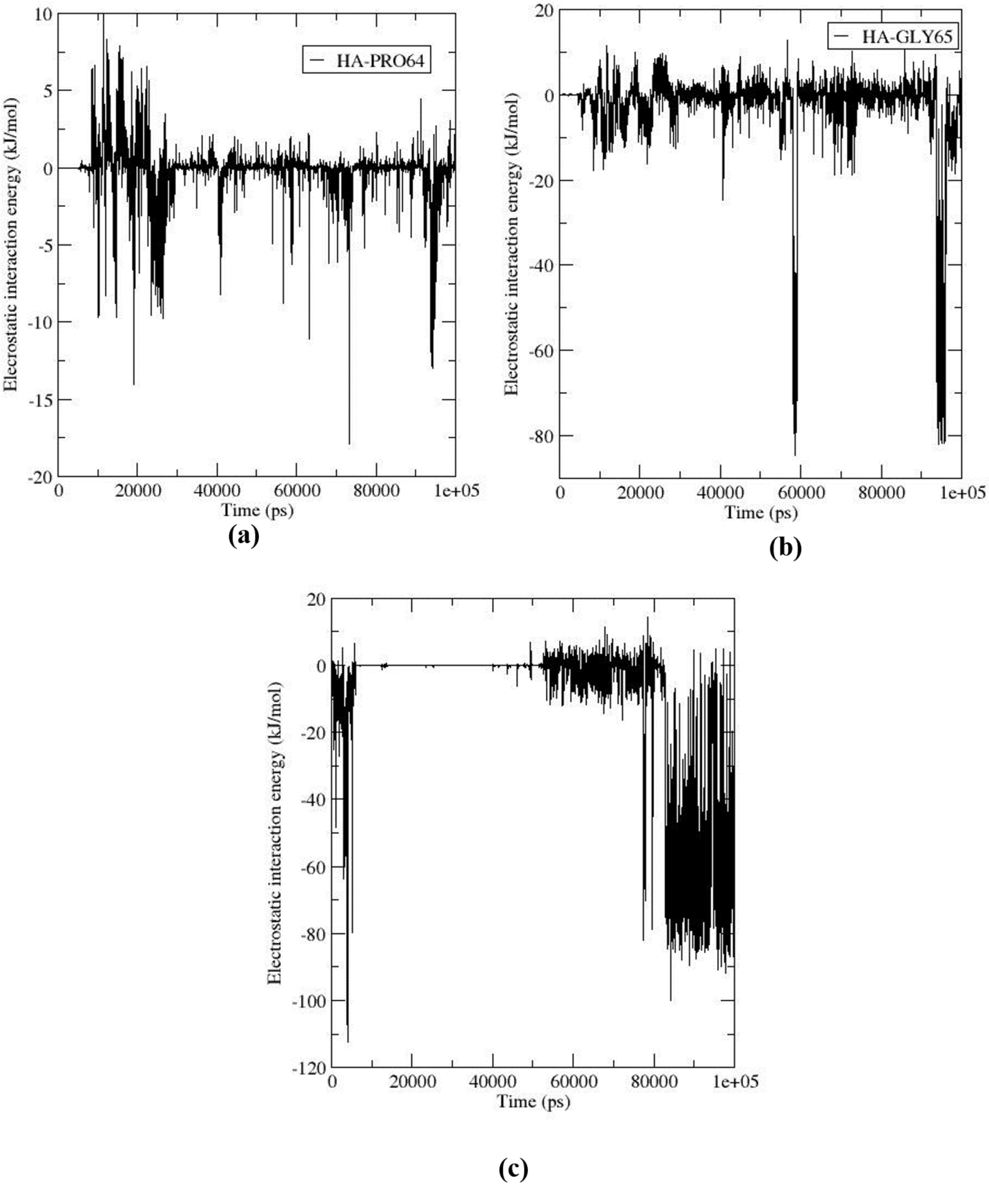


**Fig. S12:** Neutral residues interact weakly with HA surface, as evident from interaction energy of neutral residues close to surface at (a)-(b) 1.00 V/nm and (c) -1.00 V/nm.

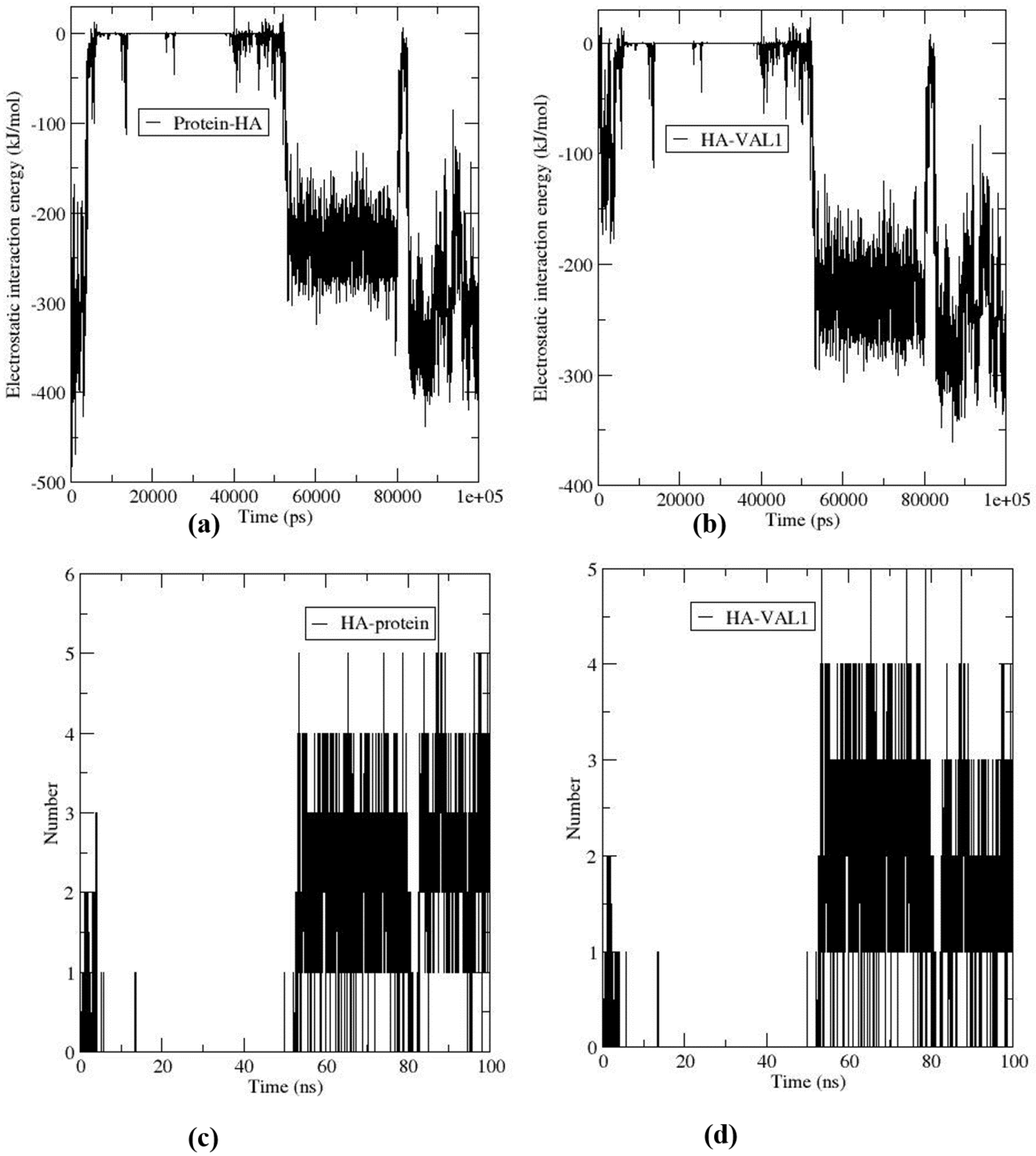


**Fig. S13:** Comparison of (a)-(b) interaction energy and (c)-(d) number of hydrogen bonds between protein-HA and VAL1-HA at electric field strength of -1.00 V/nm.

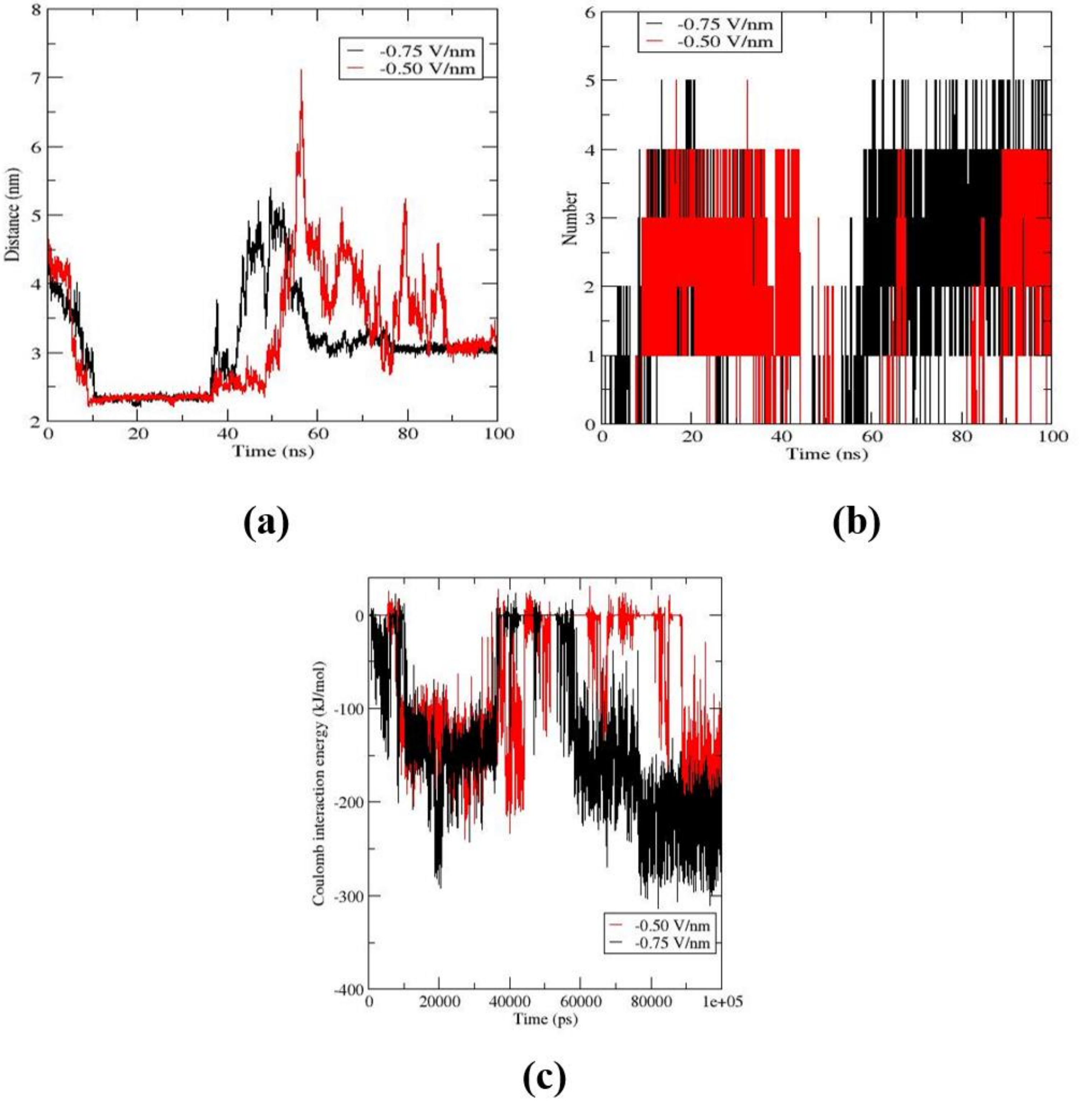


**Fig. S14:** (a) COM-COM distance (b) number of hydrogen bonds and (c) electrostatic interaction energy between Arg78 and HA at different time points.

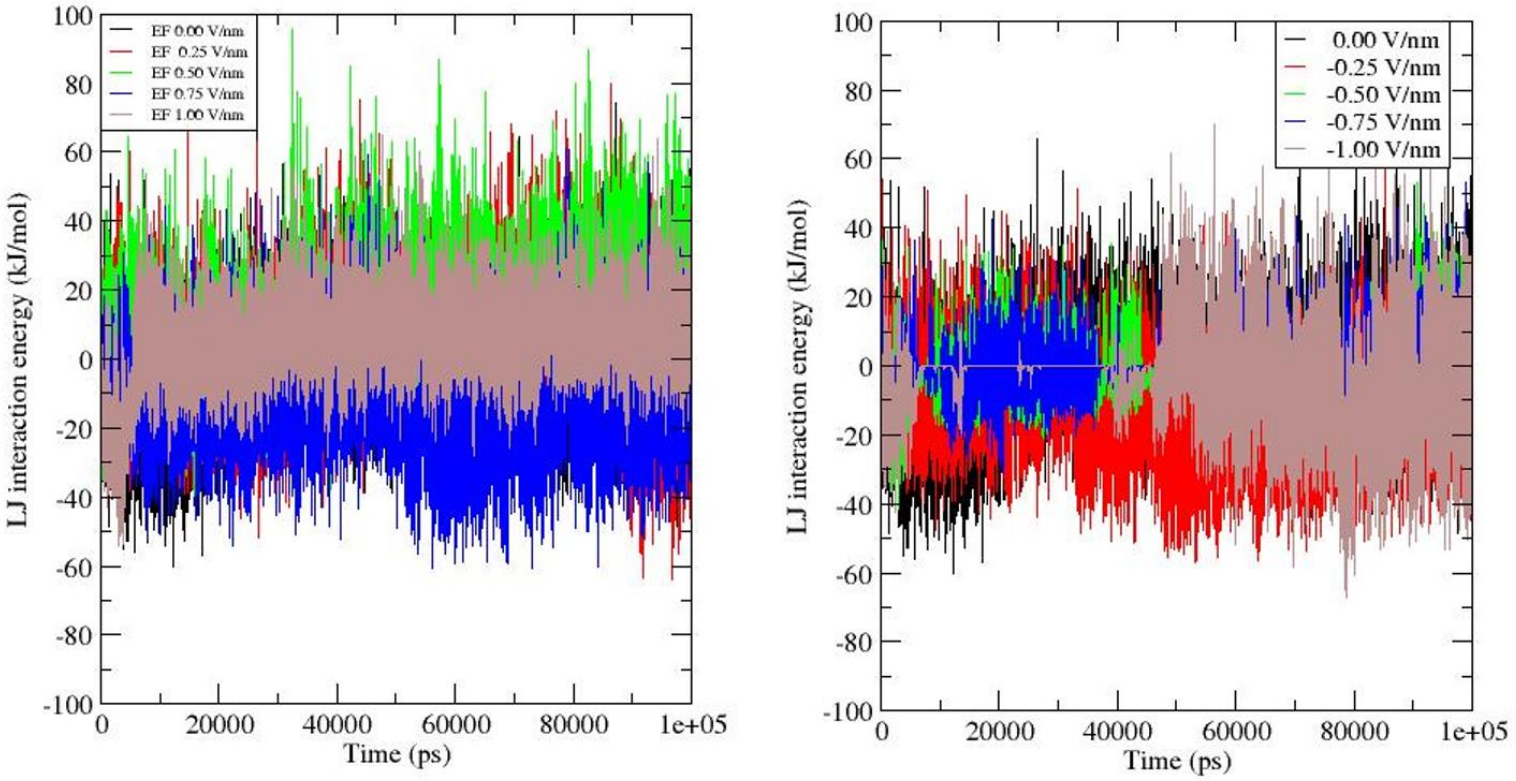


**Fig. S15:** Non-bonded interaction energy between HA and FN at different electric fields.
